# Selective Recognition, Covalent Linkage and Chemical Exchange of Artificial Cells through Engineered α-Hemolysin

**DOI:** 10.64898/2026.10.16.759538

**Authors:** Xu Han, Yuta Ishii, Kota Naito, Ayako Ijuin, Takayoshi Watanabe, Wancheng Zhang, Ako Kagawa, Asako Narita, Takeshi Yokoyama, Toshihisa Osaki, Shoji Takeuchi, Ryuji Kawano, Yoshikazu Tanaka, Tomoaki Matsuura

**Affiliations:** Department of Life Science and Technology, Institute of Science Tokyo, Tokyo 152-8550, Japan; Graduate School of Life Sciences, Tohoku University, Sendai 980-8577, Japan; Department of Biotechnology and Life Science, Tokyo University of Agriculture and Technology (TUAT), Koganei, Tokyo 184-8588, Japan; Earth-Life Science Institute (ELSI), Institute of Science Tokyo, Tokyo 152-8550, Japan; Tohoku University Advanced Research Center for Innovations in Next-Generation Medicine (INGEM), Tohoku University, Sendai 980-8573, Japan; Artificial Cell Membrane Systems Group, Kanagawa Institute of Industrial Science and Technology, Kawasaki 213-0012, Japan; Graduate School of Information Science and Technology, The University of Tokyo, Tokyo 113-8656, Japan

**Author notes:** These authors contributed equally to this work (Xu Han, Yuta Ishii). Corresponding authors YT TM.

## Abstract

Bottom-up synthetic biology aims not only to build individual artificial cells but also to connect them into higher-order systems that communicate with each other. Reported artificial cells communicate through partner recognition, physical connection, and chemical signaling that are typically implemented by separate molecular species. Here, we engineer the pore-forming protein α-hemolysin (AH) with the SpyTag–SpyCatcher (ST-SC) pair to integrate partner recognition, covalent linkage between cells, and nanopore-mediated connection of their lumens into a single protein system. Fusion of ST or SC to AH preserved heptamer assembly and pore formation, and allowed the two heptamers (AH-ST and AH-SC) to be covalently joined. Cryo-electron microscopy resolved the resulting head-to-head 14-mer assembly at 2.20 Å. Calcein leakage and single-channel recordings confirmed that the engineered pores remain functional. Mixing ST- and SC-displaying giant unilamellar vesicles (GUVs) resulted in partner-specific association between the two vesicle populations. Finally, photo-triggered release of a newly synthesized caged ethidium bromide (EtBr) in sender GUVs produced EtBr accumulation in receiver GUVs that required direct physical linkage between the two GUVs. This platform provides a modular route toward artificial-cell assemblies in which both connectivity and cargo exchange are programmable.

## Introduction

Bottom-up synthetic biology seeks to reconstruct cellular functions from simplified components and has produced diverse artificial-cell platforms based on lipid vesicles, coacervates, and other protocell-inspired systems^1–4^. As the field moves beyond individual compartments toward higher-order assemblies, communication between artificial cells has become increasingly important for constructing coordinated multicompartment systems^5–7^. In living tissues, direct cell-cell communication often couples selective physical contacts with molecular exchange through intercellular channels such as gap junctions^8^. In this sense, contact defines who to communicate, while the channel defines content of the communication.

In many of the previously reported artificial cellular systems, communication is indirect: a soluble signal is released from the sender compartment into the surrounding medium and subsequently detected by the receiver compartment^9–15^. Because the signal passes through the bulk, it is diluted and becomes available to any compartment nearby. Such communication pathway does not define a unique sender-receiver pair. In other approaches, compartments are designed to associate physically through membrane-anchored affinity pairs, DNA hybridization, or reconstituted tight-junction proteins^16–20^, while adhesion itself did not provide a connection between their lumen. More recent designs have coupled the two, pairing adhesion with pore-forming proteins or with docked synthetic nanopore, so that adhering compartments can also exchange chemicals^21–23^. In these systems, compartments are brought together by non-covalent interactions such as electrostatic and affinity-based interactions. These interactions are reversible and sensitive to solution conditions, which limits the stability of the resulting connection and makes it difficult to hold assemblies of many selectively paired populations together once formed. In artificial cellular systems, therefore, the remaining challenge is not only to integrate direct communication, but also to make the resulting connection covalent, stable, and highly modular.

α-Hemolysin (AH) is attractive for this purpose because it spontaneously inserts into lipid bilayers and forms a well-characterized heptameric nanopore that permits the passage of small molecules^24–27^. AH has been widely used to support communication between an artificial cell and the environment, between two artificial cells, and between an artificial and living cells^28–32^. However, AH itself carries no partner-recognition function, and pores residing in two apposed membranes are not connected with each other.

Here, we used giant unilamellar vesicles (GUVs) as model artificial-cell compartments and engineered AH with the SpyTag-SpyCatcher (ST-SC) system to establish an artificial-cell communication platform that integrates vesicle recognition, physical linkage, and nanopore-dependent direct molecular transport within a single protein-engineered system. ST-SC spontaneously forms an irreversible covalent isopeptide bond with high specificity and efficiency^33,34^. AH fused to ST-SC system therefore provides complementary functions: AH enables spontaneous membrane insertion and nanopore formation, whereas ST-SC specifies and covalently connects two GUVs together. We show that the engineered fusion proteins preserve pore-forming activity while enabling selective linkage between GUV populations. Furthermore, a photoactivatable molecular cargo enabled UV-triggered release and uptake in GUV assemblies. This modular strategy combines molecular recognition with transmembrane transport and establishes a framework for constructing increasingly complex communicating artificial-cell networks.

## Results

### Engineering of ST-SC-functionalized AH

To establish an artificial vesicle communication platform capable of specific molecular recognition and nanopore-mediated transport, we constructed AH proteins with either ST or SC (Figure 1a). We also developed a photocaged compound to control the transport across the nanopore. The AH was genetically fused to either ST or SC at its N- or C-terminus. For consistency with our established AH construct, the G122S/K147R background, known to improve the pore-forming activity^35^, is referred to hereafter as WT, whereas K237C denotes the same background carrying an additional K237C substitution. K273C is a mutation proposed previously that located in the position where two heptamers could form a disulfide bridge when head-to-head 14-mer is formed^36^. We first evaluated whether the fusion affected the ST-SC conjugation. Both N- and C-terminal fusion proteins were expressed using a reconstituted cell-free protein synthesis system (PURE system). Complex formation between ST and SC was detected for the C-terminal fusion, whereas little complex formation was observed for the N-terminal fusions (Figure S1). The trend was observed for both the WT and K237C constructs. All subsequent experiments were performed using the C-terminal constructs, hereafter referred to as WT-ST, WT-SC, K237C-ST, and K237C-SC, respectively.

**Figure 1.**
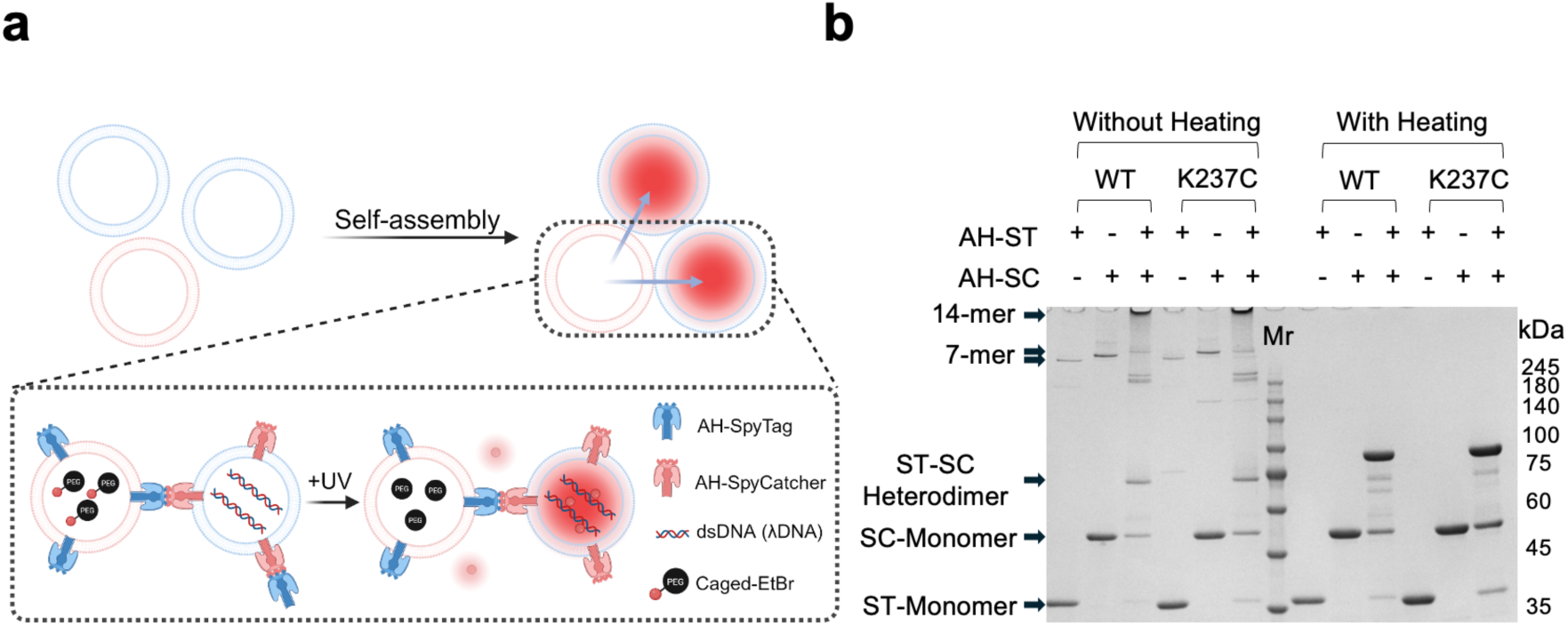
AH conjugation via ST and SC. **a)** Schematic illustration of the artificial vesicle communication system. AH was genetically fused to ST or SC to enable vesicle linkage through ST-SC conjugation. Upon UV irradiation, caged ethidium bromide (caged-EtBr) encapsulated in the sender GUV releases free EtBr, which can either reach the receiver GUV through the linked AH assembly or escape into the external medium. EtBr entering the receiver lumen binds the encapsulated DNA and generates a fluorescence signal. **b)** SDS-PAGE analysis of AH-ST and AH-SC under non-heated and heated conditions. WT and K237C constructs were analyzed individually or after equimolar mixing.

We next examined whether the ST or SC fused AH proteins retained their ability to oligomerize into heptamers. All proteins were analyzed by SDS-PAGE individually and as equimolar mixtures of ST and SC constructs (WT-ST with WT-SC, or K237C-ST with K237C-SC) under non-heated and heated conditions (Figure 1b). Because the AH heptamer is resistant to SDS unless heated (95 °C, 8 min), oligomeric species remain intact under non-heated conditions. A band corresponding to the heptamer was detected in all non-heated samples, indicating that neither fusion abolished heptamerization. An additional band of higher molecular mass was observed only in the mixed samples and was absent when AH-ST or AH-SC was analyzed alone. This species was consistent with a higher-order assembly formed by covalent ST-SC conjugation between two AH oligomers. Heating abolished both the heptameric and higher-order oligomeric bands, whereas a band corresponding to the covalent ST–SC heterodimer was retained. Heat treatment therefore disrupted the non-covalent contacts responsible for AH oligomerization while leaving the ST–SC isopeptide bond intact, and the same behavior was observed for both the WT and K237C constructs.

To verify the oligomeric states of the engineered proteins, we further analyzed the samples by size-exclusion chromatography (SEC) (Figure S2). Under non-heated conditions, the mixed samples displayed an additional early-eluting peak that was absent from either AH-ST or AH-SC alone, indicating the presence of a larger molecular assembly. Following heating, this peak disappeared, whereas the peak corresponding to the covalently linked heterodimer was retained. The SEC profiles therefore closely mirrored the SDS-PAGE results. Together, these findings demonstrate that C-terminal fusion preserves both efficient ST-SC conjugation and the intrinsic oligomerization capability of AH.

### Structure determination of head-to-head AH assemblies by cryo-EM

Previous study has modeled the head-to-head bound 14-mer^36^, while the actual structure remained unsolved. To define the structural architecture of the engineered AH assemblies, we performed cryo-EM analysis of the WT and K237C AH-ST/AH-SC complexes. Purified WT-ST and WT-SC, or K237C-ST and K237C-SC, were mixed at a 1:1 molar ratio to allow ST-SC-mediated complex formation before cryo-EM analysis. In both samples, we observed abundant particles consistent with head-to-head 14-mer assemblies, using transmission electron microscopy (TEM), formed by the association of two AH heptamers (Figure S3, a). Subsequent single-particle analysis enabled reconstruction of the WT and K237C 14-mer assemblies at a resolution of 2.20 Å.

In both the WT (PDB 8JX2) and K237C (PDB 8JX3) assemblies, the overall structure of each AH heptamer closely resembled previously reported AH structures^24^, and no substantial structural differences between the two variants. In both structures, the ST and SC regions were not resolved, suggesting the with conformational flexibility of these regions. Consistent with this interpretation, Gaussian-filtered maps revealed weak and unresolvable densities at seven positions near the interface between the two opposing heptamers (Figure S3, b). These findings indicate that while the ST-SC linkages remain flexible, the AH 7-mers associate with each other to form a defined structure regardless of the presence of the Cys residue.

Comparison of the WT and K237C structures revealed a distinct difference at the inter-heptamer interface (Figure 2b, c). In the K237C assembly, Cys237 residues from opposing heptamers formed disulfide bonds at seven equivalent positions around the interface (Figure 2c). Structural superposition of the WT and K237C assemblies showed that the upper heptamer in the K237C structure was rotated by 8.9° relative to that in the WT structure (Figure 2b). This rotation appears to accommodate disulfide bond formation between opposing Cys237 residues. In contrast, the corresponding Lys237 residues in the WT structure were separated by 14.9 Å (Nζ-Nζ) and did not form equivalent inter-heptamer contacts (Figure 2c). Thus, although both WT and K237C form head-to-head 14-mer assemblies, the K237C substitution introduces additional disulfide-mediated contacts that alter the relative orientation of the two heptamers.

**Figure 2.**
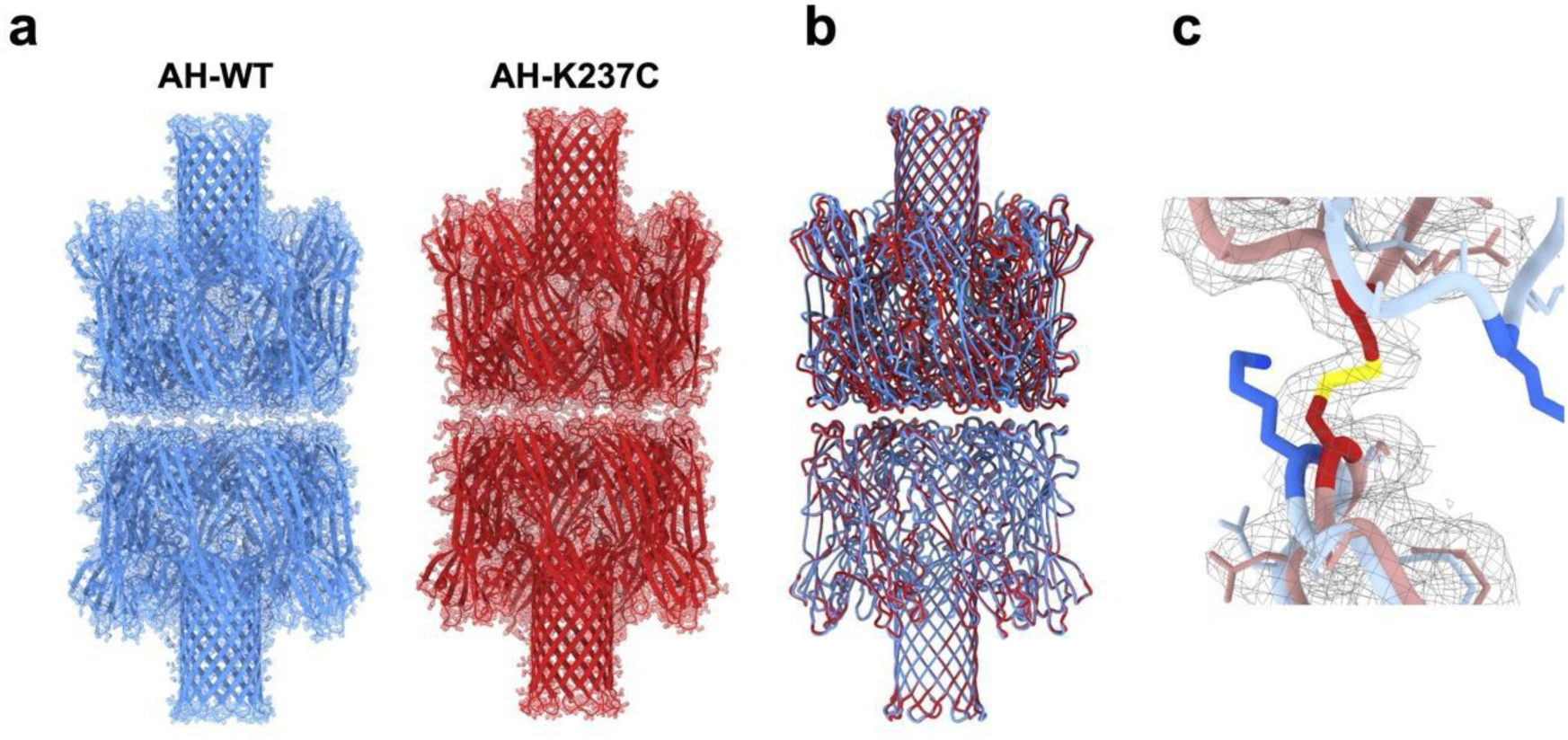
Cryo-EM structures of WT and K237C AH 14-mer assemblies. **a)** Cryo-EM structures of the WT and K237C AH 14-mer assemblies. Each 14-mer formed by head-to-head association of two AH heptamers. Space-filling and ribbon representations were overlaid. **b)** Overlay of WT and K237C structures. The bottom 7-mers were superimposed completely, while the upper 7-mer of K237C was rotated by 8.9° relative to that of WT. **c)** Close-up view of residue 237 at the inter-heptamer interface. WT (blue) and K237C (red) structures were shown superimposed; only residue 237 was highlighted in saturated colors, with surrounding residues shown in pale colors. The grey mesh represented the cryo-EM density of K237C (EMD-36689, contoured at 0.108), shown only around the K237C model; the corresponding WT density (EMD-36688) was not displayed for clarity. Sulfur atoms of Cys237 were shown in yellow; the continuous density spanning the two sulfur atoms supported the presence of the disulfide bond. Opposing Lys237 residues in WT remain separated (Nζ-Nζ), 14.9 Å), whereas opposing Cys237 residues in K237C formed an inter-heptamer disulfide bond (Sγ–Sγ, 2.0 Å).

### Pore-forming assay of engineered AH fusion proteins

We next examined whether fusion of ST or SC affected the pore-forming function of AH. Pore activity was evaluated by a calcein leakage assay, in which calcein (0.6 kDa) and ovalbumin–Alexa Fluor 647 (OA647, ∼45 kDa) were co-encapsulated in GUVs. OA647 served as a non-permeable internal reference for normalization. Calcein retention was quantified as the geometric mean of the calcein/OA647 fluorescence measured by flow cytometry (FCM), with lower values (relative calcein concentration) indicating greater leakage.

All four AH fusion proteins (WT-ST, WT-SC, K237C-ST, and K237C-SC) induced concentration-dependent calcein leakage (Figure 3a). Representative two-dimensional FCM plots of WT-ST showed a pronounced concentration dependence, whereas changes associated with longer incubation times were comparatively smaller (Figure S6). Under all protein concentrations tested, the calcein/OA647 value approached a plateau after approximately 5 h of incubation (Figure S7).

**Figure 3.**
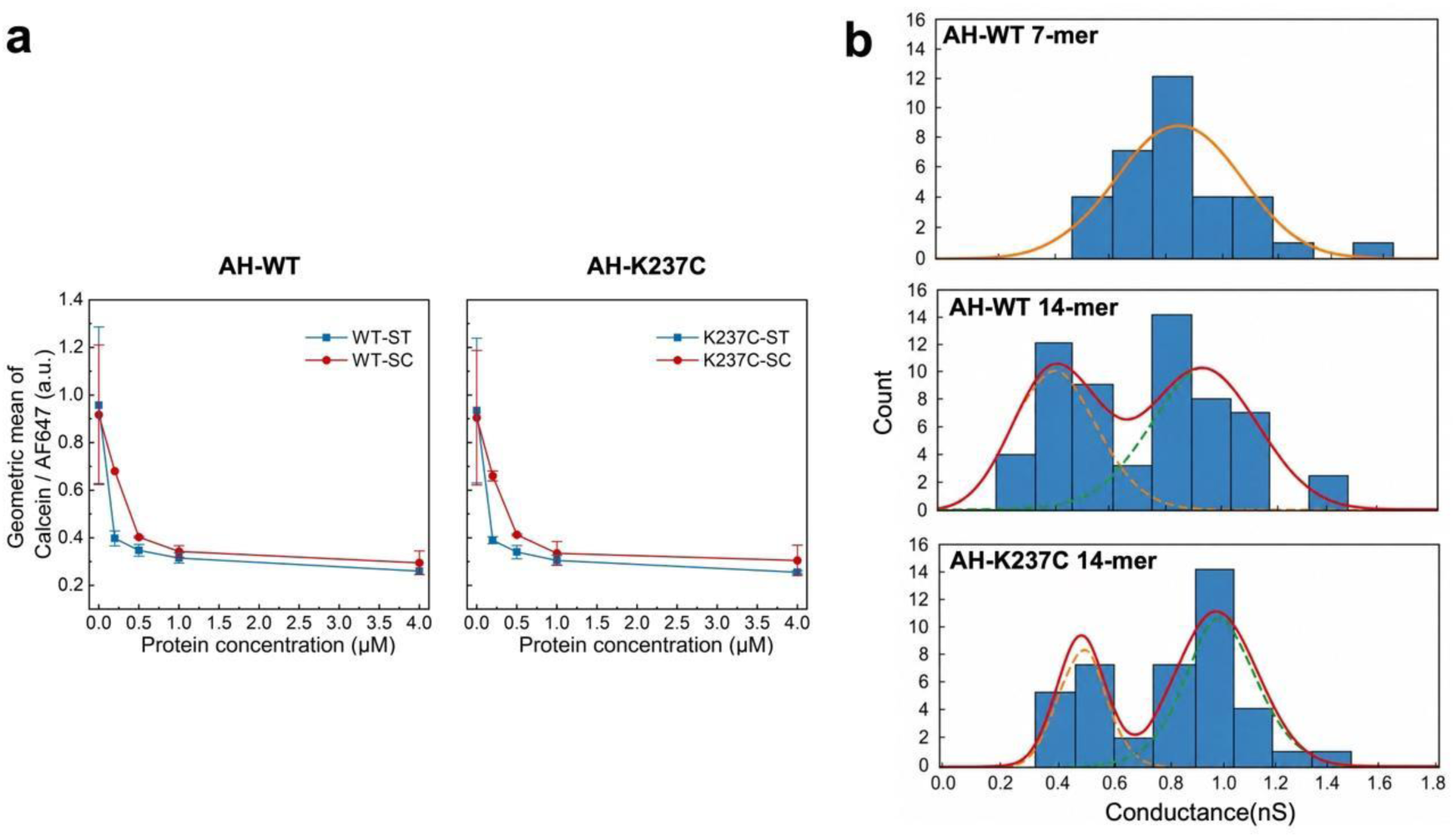
Pore-forming activity and single-channel conductance of engineered AH fusion proteins. **a)** Calcein leakage assay of engineered AH fusion proteins. GUVs co-encapsulating calcein and OA647 were incubated with AH fusion proteins for 5 h. Leakage was quantified as the geometric mean of the calcein/OA647 fluorescence measured by FCM. Data are the mean ± SD from two independent GUV preparations, each measured in duplicate. **b)** Single-channel conductance analysis of engineered AH nanopores measured using DIBs. Conductance histograms were fitted using Gaussian models. The WT 7-mer was adequately described by a single conductance population, whereas the WT and K237C 14-mers samples exhibited bimodal conductance distributions consistent with two conductive pore species.

To assess the functionality of the engineered nanopores more quantitatively, we measured single-channel conductance using droplet interface bilayers (DIBs) (Figure 3b). The WT 7-mers exhibited a relatively narrow conductance distribution that was adequately described by a single Gaussian population, with a mean conductance of 0.85 nS in 1 M KCl. This value closely matches the 0.857 nS reported for the heptameric AH pore under comparable ionic conditions^36^, indicating that C-terminal fusion of ST does not alter the conductive properties of the pore.

In contrast, the WT and K237C 14-mers each exhibited bimodal conductance distributions that were resolved into two populations by Gaussian mixture analysis. For WT 14-mers, the populations were centered at approximately 0.38 nS and 0.93 nS, and for K237C 14-mers at approximately 0.47 nS and 0.99 nS. The higher-conductance population in each sample overlapped with the range observed for the 7-mers and is therefore assigned to unconverted heptamers, consistent with the incomplete conversion to the covalently linked assembly indicated by SDS-PAGE and SEC (Figure 1b and Figure S2). The lower-conductance population is instead consistent with the reduced conductance previously reported^36^ for the head-to-head pair of AH heptamers, in which one β-barrel spans the bilayer while the second remains in solution, adding a second pore in series (0.57 nS, compared with 0.857 nS for the heptamer).

Together, the leakage and single-channel measurements show that the ST- and SC-fused AH constructs retain pore-forming activity and that the covalently associated assemblies remain electrically conductive.

### Selective vesicle recognition and linkage mediated by ST-SC interactions

We next investigated whether ST and SC sequences could mediate selective association between GUV populations. Two GUV populations, distinguished by encapsulated GFP or OA647, were separately incubated with AH-ST or AH-SC, mixed at a 1:1 ratio, and analyzed by FCM. FCM events were divided into four quadrants: Q1, OA647-positive/GFP-negative; Q2, OA647-positive/GFP-positive; Q3, GFP-positive/OA647-negative; and Q4, double-negative (Figure 4a). The Q2 fraction (black-boxed region in Figure 4a) was used as an operational measure of association between the two fluorescently distinct GUV populations. In the absence of AH proteins, only a small fraction of events was detected within Q2, indicating minimal spontaneous vesicle association. Similarly, the SpyTag/SpyTag (ST/ST) and SpyCatcher/SpyCatcher (SC/SC) control groups exhibited only low levels of double-positive events, indicating that nonspecific vesicle association remained limited. In contrast, the SpyTag/SpyCatcher (ST/SC) group showed a substantial increase in the Q2 fraction (50 to 60%, on average), indicating that association between the two GUV populations was strongly enhanced by complementary ST-SC recognition (Figure 4a).

**Figure 4.**
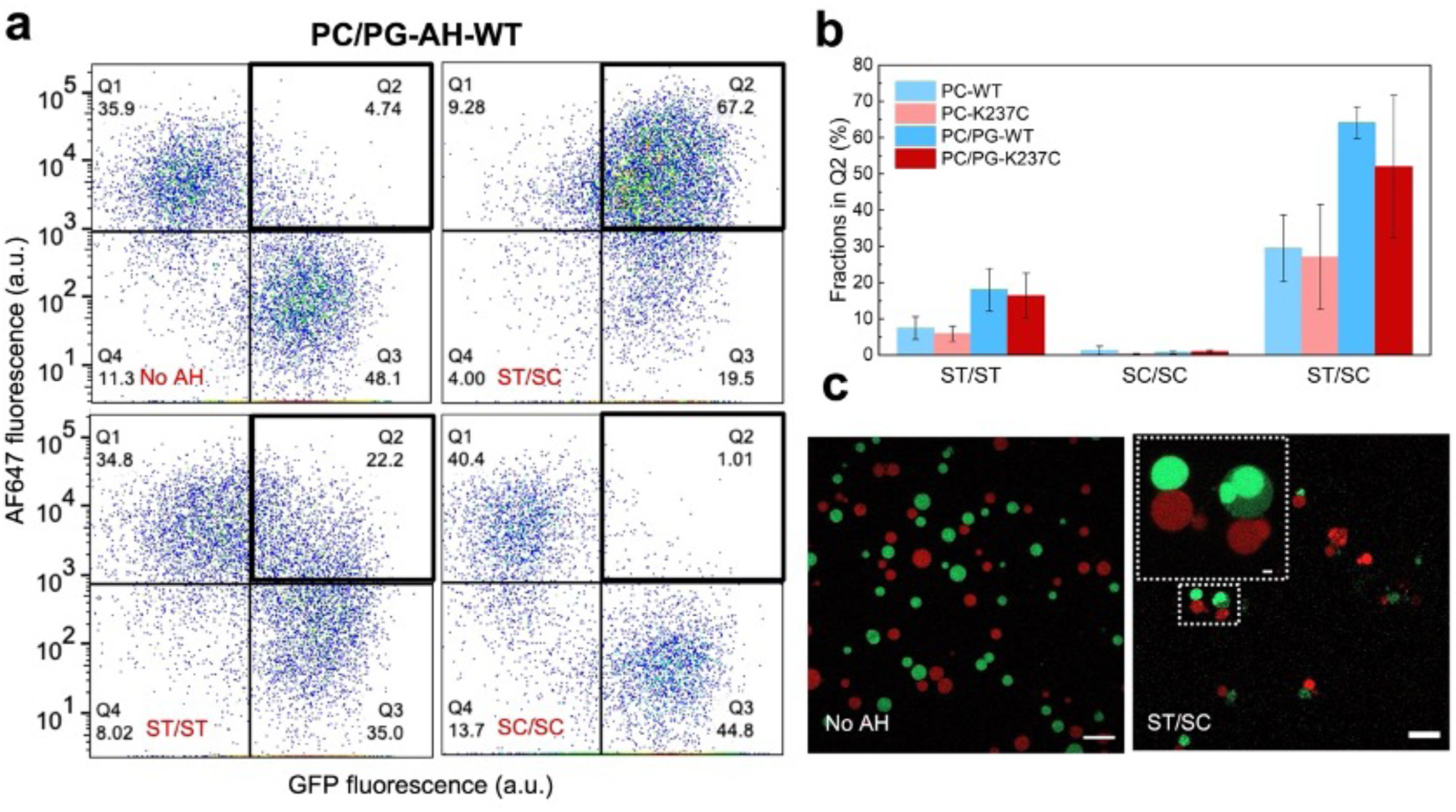
Flow cytometric and microscopic characterization of vesicle linkage. **a)** Representative FCM plots for the WT constructs under the optimized PC/PG lipid composition. Q1, OA647-positive/GFP-negative; Q2, OA647-positive/GFP-positive; Q3, GFP-positive/OA647-negative; Q4, double-negative. **b)** Quantification of vesicle association efficiency measured as the fraction of Q2 events for WT and K237C constructs prepared under PC or PC/PG lipid compositions. ST/ST, SpyTag/SpyTag; SC/SC, SpyCatcher/SpyCatcher; ST/SC, SpyTag/SpyCatcher. For PC conditions, data are mean ± SD from two independent GUV preparations. For PC/PG conditions, data are mean ± SD from three independent GUV preparations. **c)** Confocal fluorescence images of GUV populations. Left, GUVs without AH proteins. Right, mixture of two GUV populations functionalized with AH-ST and AH-SC (1 μM each), respectively. Green, SNAP-GFP; red, mCherry. Inset shows an enlarged view of the boxed region. Scale bars: 20 μm; inset, 2 μm.

During optimization of the GUV linkage assay, we found that incorporation of the anionic lipid POPG substantially increased vesicle-linkage efficiency (Figure S9). Under the PC/PG condition, the ST/SC fraction increased from approximately 28% to 58%, while the ST/ST control also rose from approximately 7% to 17%; the SC/SC control remained unchanged at ∼1% (Figure 4b). The ST/SC group nevertheless remained approximately three-fold higher than the ST/ST control, indicating that specific ST-SC recognition remained the dominant interaction under the optimized composition. To further assess whether Q2 events were consistent with physically associated GUVs, we compared the forward-scatter distributions of the gated populations (Figure S10). Within the ST/SC sample, Q2 events exhibited higher forward scatter than those in Q1 and Q3 populations, consistent with the larger apparent size expected for associated vesicles or vesicle clusters.

Furthermore, GUV populations were directly visualized by confocal fluorescence microscopy (Figure 4c). In the absence of AH proteins, the two GUV populations remained dispersed (Figure 4c, left). In contrast, GUVs functionalized with complementary AH-ST and AH-SC were frequently observed in direct contact, forming small mixed clusters (Figure 4c, right). Together with the FCM data, these observations support ST-SC-mediated physical association between the two independently prepared GUV populations.

### Photo-controlled cargo release and uptake in linked GUV populations

To achieve temporally controlled molecular exchange between linked vesicles, we synthesized a photoactivatable caged-ethidium bromide (caged-EtBr) that releases free EtBr upon UV irradiation (Figure 1a, 5a and S11). The caged-EtBr was conjugated to mPEG5000 to prevent passage of the compound through the AH pore before photoactivation. We first validated its photoactivation and DNA-dependent fluorescence enhancement in bulk solution (Figure S12) and then in GUVs (Figure 5b). GUVs containing caged-EtBr together with λDNA exhibited low EtBr fluorescence before UV irradiation and a pronounced fluorescence after UV irradiation. GUVs containing caged-EtBr alone also displayed slight increased fluorescence after UV irradiation, reflecting the intrinsic fluorescence of free and uncaged EtBr, while the fluorescence intensity remained lower than that observed in the presence of λDNA (Figure 5b). GUVs containing λDNA alone showed no detectable fluorescence before or after UV irradiation. Furthermore, whether the uncaged-EtBr could permeate through AH pores was analyzed by externally supplying caged-EtBr, uncaging followed by the analysis of the intravesicular EtBr signal. Strong fluorescence was observed only when AH was present (Figure S13), indicating that uncaging induces an increase in the EtBr fluorescence. Together, these results showed that caged-EtBr/λDNA as a suitable reporter pair for detecting UV-triggered EtBr release and accumulation.

**Figure 5.**
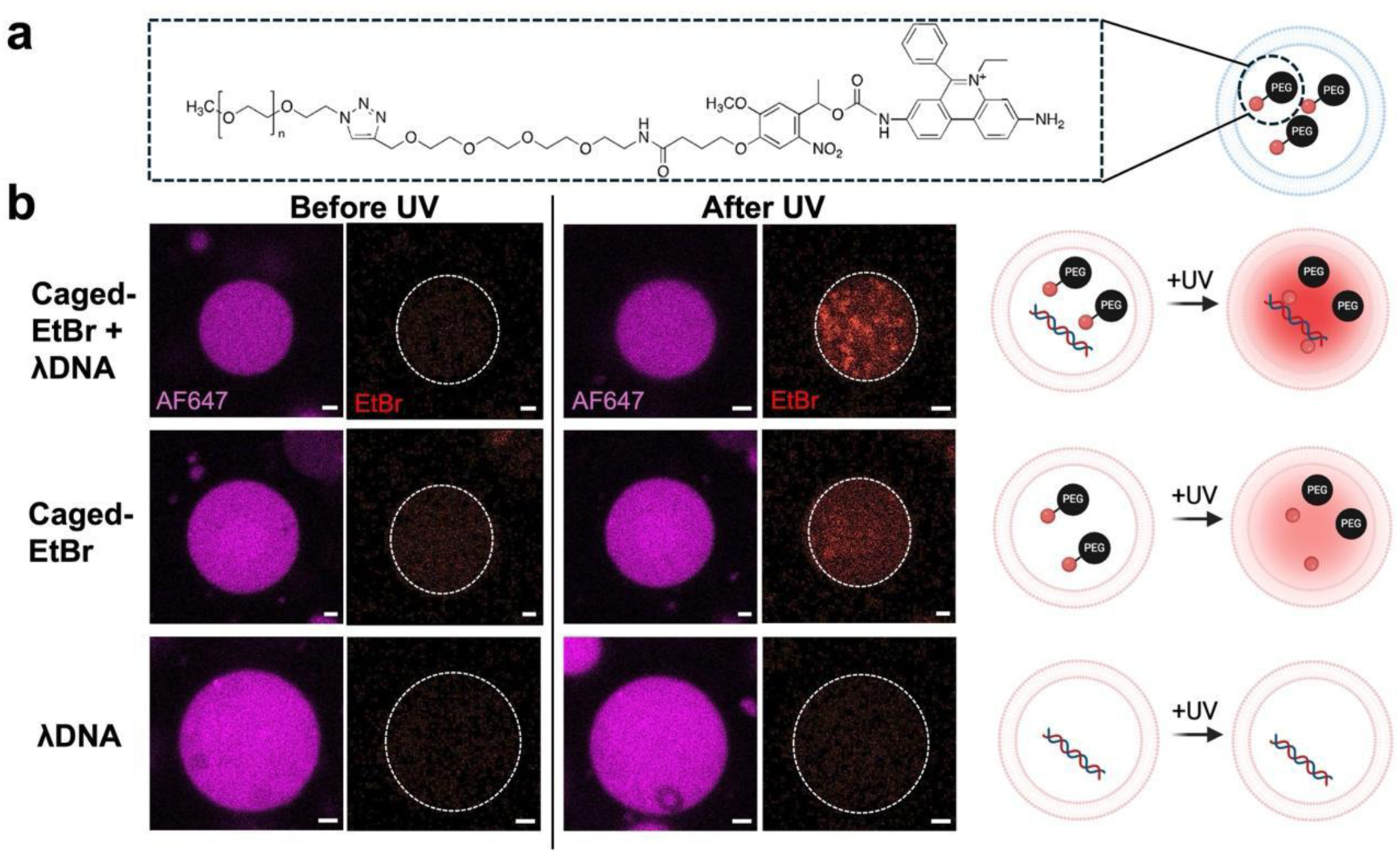
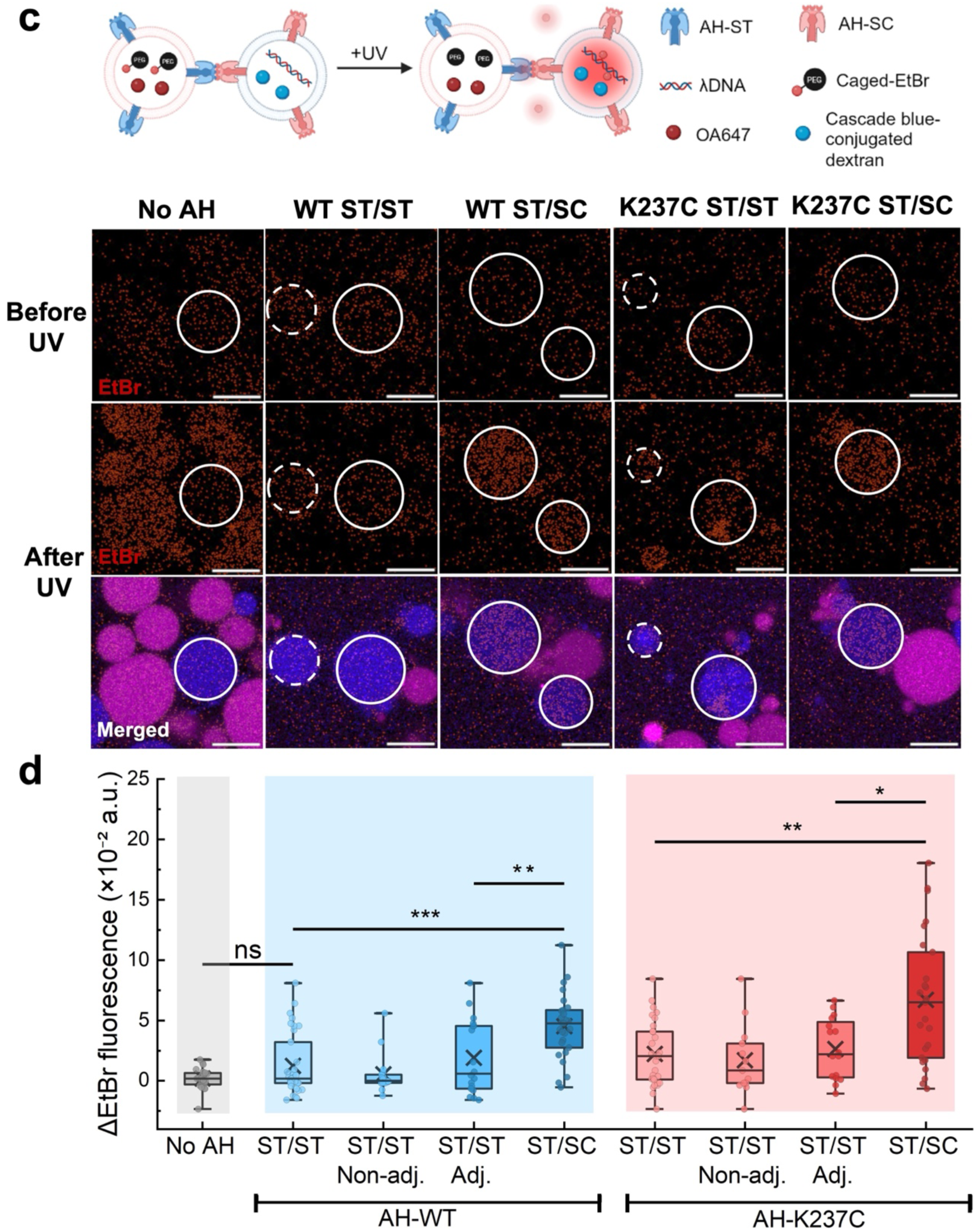
UV-triggered EtBr release and accumulation in linked receiver GUV population. **a)** Chemical structure of caged-EtBr and schematic illustration of UV-triggered release of free EtBr from sender vesicles. **b)** Validation of caged-EtBr photoactivation. Representative confocal fluorescence images of GUVs encapsulating caged-EtBr and λDNA, caged-EtBr alone, or λDNA alone before and after UV irradiation. Schematics illustrating the expected fluorescence response are shown on the right. Scale bars: 2 μm. **c)** Top, schematic of the transport experiment: sender GUVs encapsulate caged-EtBr and OA647, receiver GUVs encapsulate λDNA and cascade blue-conjugated dextran. Bottom, representative confocal images of receiver GUVs for the five conditions. The EtBr channel is shown before (top) and after (middle) UV irradiation; the merged image (bottom) additionally shows OA647 (sender) and cascade blue (receiver). Circles mark the receiver GUVs analyzed; in the ST/ST conditions, solid circles represent receivers adjacent to a sender vesicle and dashed circles represent non-adjacent receivers. Brightness and contrast were adjusted identically for all images for better visualization. Scale bars: 10 μm. **d)** Per-vesicle EtBr fluorescence increase *(ΔF = F_after_ − F_before_)*. Groups: 1, no AH; 2 - 5, AH-WT (2, ST/ST pooled; 3, ST/ST non-adjacent; 4, ST/ST adjacent; 5, ST/SC); 6 - 9, AH-K237C (6, ST/ST pooled; 7, ST/ST non-adjacent; 8, ST/ST adjacent; 9, ST/SC). The pooled ST/ST groups (2 and 6) are the union of the corresponding non-adjacent and adjacent groups and are not independent of them. Boxes indicate the median and interquartile range, whiskers the full data range, crosses the mean; individual vesicles are overlaid as points. *n* = 23, 35, 17, 18, 27, 29, 13, 16 and 25 GUVs for groups 1 - 9. Statistically significant differences are indicated based on two-sided Mann–Whitney *U* tests: 1 vs 2, *p* = 0.622; 2 vs 5, *p* < 0.0001; 3 vs 4, *p* = 0.488; 4 vs 5, *p* = 0.0071; 6 vs 9, *p* = 0.00169; 7 vs 8, *p* = 0.254; 8 vs 9, *p* = 0.0168. \**p* < 0.05, \*\**p* < 0.01, \*\*\**p* < 0.001; n.s., not significant.

Finally, we examined whether UV irradiation produces a detectable EtBr signal in the linked receiver vesicles, and whether that signal depends on ST/SC linkage. Sender GUVs encapsulated caged-EtBr and OA647, whereas receiver GUVs encapsulated λDNA and cascade blue-conjugated dextran (Figure 5c). The two populations were functionalized with the indicated AH constructs, mixed, placed under the microscopy, and irradiated by the UV light. EtBr fluorescence of individual receiver GUVs was quantified by confocal microscopy before and after irradiation, and the fluorescence increase (Δ*F* = *F*_after_ − *F*_before_) was used as the readout (Figure 5c). In both the WT and K237C backgrounds, ST/SC receiver GUVs showed greater EtBr fluorescence increases than the ST/ST controls indicating the direct EtBr transport through head-to-head bound AH. Note that AH could be present not at the interface between the GUVs, which could lead to the leakage of EtBr, while the signal increase in the receiver GUVs indicate the presence of direct transport through the 14-mer. More quantitative analysis is described in the Discussion section.

To further confirm the direct transport of uncaged EtBr, ST/ST population was further analyzed. Because ST/SC receivers were physically associated with sender GUVs, the pooled ST/ST population does not provide a geometry-matched control for vesicle proximity. Confocal images occasionally revealed apparent contacts between sender and receiver GUVs even in the ST/ST samples. We thus designated the receiver GUVs whose membrane were in direct apparent contact with a sender vesicle in the same focal plane as “adjacent”, and the others as “non-adjacent GUVs”. Representative images of the two classes are indicated by solid and dashed circles, respectively in Figure 5c. Adjacent and non-adjacent receiver GUVs for ST/ST sample did not differ significantly, indicating that physical proximity alone was not associated with increased EtBr accumulation (Figure 5d). Linked ST/SC receivers nevertheless accumulated significantly more EtBr than adjacent ST/ST receivers. These results support a specific contribution of ST-SC-mediated linkage on the fluorescence increase.

## Discussion

In this study, we integrated the ST-SC pair with the pore-forming protein AH to embed selective molecular recognition, vesicle association, and direct transport function to a single engineered protein. Cryo-EM showed that ST-SC conjugation yields a defined head-to-head AH 14-mer composed of two opposing heptamers. Whereas head-to-head dimerization driven by Cys237 alone yielded a minor species (<5%) when expressed in *E. coli* and that had to be isolated by preparative gel extraction^36^, the ST-SC pair provides a specific recognition and covalent linkage module, and the head-to-head assembly is obtained directly on mixing with an efficiency of ∼60% (Figure 1b), much higher than that with Cys237 alone. Although K237C introduced additional inter-heptamer disulfide bonds and altered the orientation of the two heptamers relative to that without K237C (Figure 2 c), the functional assays tested revealed no difference between WT and K237C. Because AH-WT cannot form inter-heptamer disulfides yet behaved indistinguishably from K237C in every assay, the disulfides are not required: isopeptide bonds derived from ST-SC already close the interface covalently, therefore the disulfides represent a second, redundant constraint.

The photo-controlled EtBr release experiments showed that ST-SC-mediated linkage enables direct EtBr transport from physically connected sender to receiver GUVs, with receiver signals in the ST/SC groups significantly higher than in the ST/ST control. Nevertheless, receiver EtBr signals appeared lower than in control GUVs in which EtBr and λDNA were co-encapsulated at the same initial concentrations (Figure 5b, top), indicating that direct transfer does not account for all released EtBr.

The AH nanopores on GUV membrane are located at the solvent-exposed surface and not necessarily at the GUV-GUV contact interface accounts for this. According to the lipid and AH concentrations, and assuming a GUV with 10 µm in diameter and the contact area as a flat circular disk with a contact diameter of around 2 μm (Figure S14), approximately 5 × 10⁵ AH heptamers are likely present for each GUV (Note S1). When assuming an equal distribution across the membrane, only a small fraction (1%) of these is present at the contact area, and 99% at the exposed area. If uncaged EtBr is transported equally through all pores, most of the EtBr is lost and thus not detectable in the receiver GUVs. Nevertheless, EtBr transport to the receiver GUVs was observed (Figure 5c, d). There should be a mechanism for ST-SC-mediated linkage to promote the local enrichment of AH 14-mers at the contact interface. Individual AH heptamers can diffuse laterally on the membrane, while once linked as head-to-head 14-mers at the contact area, they cannot diffuse away from the contact zone (Figure S15). This is because one of the heptamer of the 14-mer, when present in the solvent-exposed area, will have the hydrophobic transmembrane domain exposed to aqueous phase, which is energetically unfavorable. Consequently, covalent linkage may favor the retention and the local accumulation of AH 14-mers at the contact interface relative to the exposed area, facilitating direct EtBr transfer between connected GUVs. Although direct evidence for such enrichment is currently lacking, this mechanism provides a plausible explanation for the observed ST-SC dependent signal enrichment of EtBr transfer in receiver GUVs.

Indeed our estimation indicates that GUV-GUV contact interface has sufficient capacity to accommodate a higher local density of AH heptamers (Note S2).

An important feature of the present design (Figure 1a) is its modularity. Molecular recognition is genetically encoded through ST or SC fusion, rather than chemical modification of the vesicle membrane. The recognition module could therefore, in principle, be exchanged for other orthogonal Tag-Catcher pairs. In addition to the SpyTag-SpyCatcher pairs, SnoopTag-SnoopCatcher and DogTag-DogCatcher provide mutually selective recognition channels that could be assigned to different GUV subpopulations^37,38^. Incorporating multiple orthogonal pairs into one system could enable selective connectivity among defined artificial-cell populations and provide a route toward multiplexed communication within more organized artificial-cell networks. The chemical transport module (in our case AH) may likewise be varied. Because different pore-forming proteins differ in their effective pore size and transport behavior, the choice of transport module may influence the size range of cargo that can be exchanged^39^. Therefore, combining orthogonal partner recognition with selective transport modules could, in theory, allow better control and programming the modularity of the cell-cell interaction at the protein level.

In conclusion, we have shown that selective molecular recognition, stable vesicle linkage, and externally triggered chemical exchange can be encoded in a single engineered protein architecture. This platform could provide a basis for constructing selectively linked vesicle networks in which chemical exchange can be externally triggered. Extension to multiple recognition and transport modules may ultimately support the assembly of more complex multicompartment systems with selective communication pathways.

## Materials and Methods

### Lipids

1-Palmitoyl-2-oleoyl-*sn*-glycero-3-phosphocholine (POPC), 1-palmitoyl-2-oleoyl-*sn*-glycero-3-phospho-(1′-*rac*-glycerol) sodium salt (POPG), cholesterol, 1,2-distearoyl-*sn*-glycero-3-phosphoethanolamine-*N*-[methoxy(polyethylene glycol)-2000] ammonium salt (DSPE-PEG2k), 1,2-distearoyl-*sn*-glycero-3-phosphoethanolamine-*N*-[biotinyl(polyethylene glycol)-2000] ammonium salt (biotin-DSPE-PEG2k) and 1,2-diphytanoyl-sn-glycero-3-phosphocholine (DPhPC) were purchased from Avanti Polar Lipids (Alabaster, AL, USA).

### Plasmid construction

pIVEX2.3d-AH-6CHis-2fix encoding α-hemolysin (AH) with G122S/K147R mutations was used as a template to introduce K237C substitution and prepare plasmid pIVEX2.3d-AH-6CHis-2fix-K237C. In brief, site-directed mutagenesis was performed using primers K237C-FP and K237C-RP, and KOD-Plus-Neo (TOYOBO Co., Ltd., Osaka, Japan) for inverse PCR.

Two plasmids pIVEX2.3d-AH-6CHis-2fix and pIVEX2.3d-AH-6CHis-2fix-K237C were used as templates to obtain constructs containing SpyTag003 or Catcher003 at either the N- or C-terminus of AH. SpyTag003 sequence was added to the N and C terminus of AH using the Q5 Site-Directed Mutagenesis Kit (New England Biolabs, Ipswich, MA, USA) according to the manufacturer’s instructions. Primers AHST-NF/AHST-NR and AHST-CF/AHST-CR were used to introduce the tag sequence to the N- and C-terminal, respectively. SpyCatcher003 sequence was added to the N- and C-terminus of AH using Gibson assembly kit (New England Biolabs, Ipswich, MA, USA) according to the manufacturer’s instructions. Synthetic DNA encoding SpyCatcher003 sequence (Table S1) was obtained from Eurofins Genomics, PCR amplified by SC3-GS-F/SC3-GS-R or GS-SC3-F/GS-SC3-R to introduce at the SpyCatcher003 to the N or C terminus, respectively, and used for Gibson assembly. Two plasmids pIVEX2.3d-AH-6CHis-2fix and pIVEX2.3d-AH-6CHis-2fix-K237C were amplified using AH-NF/AH-NR or AH-CF/AH-CR to introduce SpyCatcher003 to the N or C terminus, respectively, and used for Gibson assembly. Finally, pIVEX2.3d-AH-6CHis-2fix-SpyTag003-N, -2fix-SpyTag003-C, -2fix-SpyCatcher003-N, -2fix-SpyCatcher003-C, and their corresponding K237C mutants were obtained.

Because the pIVEX2.3d constructs were used for cell-free protein synthesis, the C-terminal AH-ST and AH-SC variants intended for recombinant expression in *E. coli* were subcloned into pET28b by Gibson assembly. DNA fragments were obtained using 2fix-to-pET28b-F/R primers to amplify the portion encoding AH-ST/SC from the corresponding plasmids. pET28b-F/R primers were used to amplify pET28b portion of the final constructs. All primers were synthesized by Eurofins genetics and plasmid sequences were confirmed by Sanger sequencing.

### Protein expression and purification

The four C-terminal AH fusion constructs (WT-ST, WT-SC, K237C-ST, and K237C-SC) in pET28b were transformed into *E. coli* BL21(DE3)-pLysS. Cells were grown in TB medium at 37 °C to an OD600 of 0.7–1.0. Protein expression was induced by addition of IPTG to a final concentration of 0.5 mM, followed by incubation at 37 °C for 5 h. Cells were harvested and the recombinant proteins were purified by immobilized metal-affinity chromatography (IMAC). The pellet was suspended in equilibration buffer (50 mM Tris-HCl [pH 8.0], 300 mM NaCl, 0.05 vol% Tween20, 10% glycerol, 7 mM 2-mercaptoethanol, 7 M urea), and the soluble fraction was mixed with TALON^®^ metal-affinity resin equilibrated with equilibration buffer. The resin was sequentially washed with equilibration buffer containing 20 mM imidazole and then with equilibration buffer containing 20 mM imidazole and 1 M NaCl. Finally, proteins were eluted with equilibration buffer containing 300 mM imidazole. The eluate was dialyzed first with buffer A (20 mM HEPES-KOH [pH 7.6], 300 mM NaCl, 5 mM 2-mercaptoethanol, 3.5 M urea) at 4°C for 1 day followed by buffer A without urea at 4°C for 2 days. After dialysis, Protein concentrations were determined from absorbance at 280 nm using calculated extinction coefficients and concentrated to more than 5 mg/mL.

### Evaluation of N- and C-terminal ST/SC fusion constructs

AH variants were synthesized in 20 µL reaction using a reconstituted cell-free protein synthesis system (PUREfrex 1.0) according to the manufacturer’s instructions. The template DNAs for the PURE system reactions were PCR-amplified using KOD-Plus-Neo, with the corresponding pIVEX plasmids (Table S2) as templates and T7F/T7R as primers. Amplification products were purified with NucleoSpin Gel and PCR Clean-up kit (Macherey-Nagel, Düren, Germany), and the concentration was measured from the absorbance at 260 nm obtained with a NanoDrop 2000 (Thermo Fisher Scientific, Waltham, MA, USA). Reactions were performed at 37°C for 4 h at a DNA concentration of 300 nM.

To evaluate covalent complex formation between AH-ST and AH-SC, 6 μL each of cell-free synthesized products were mixed and incubated at 37°C for 3 h, followed by SDS-PAGE and Western blotting using Anti-6His-HRP antibody (MBL). Luminescence detection was performed with Amersham ECL Western blot detecting reagent and imaging was performed with Typhoon FLA 9500 imaging system (GE).

### SEC analysis

SEC analysis was performed using a Superose 6 Increase 10/300 GL column (Cytiva, Marlborough, MA, USA) at 25 °C on Shimadzu HPLC LC-20A system (Shimadzu Corporation, Kyoto, Japan). The column was equilibrated with SEC buffer (20 mM HEPES-KOH (pH 7.6), 300 mM NaCl, and 0.5% (w/v) SDS). Proteins were eluted at a flow rate of 0.5 mL min⁻¹, and chromatograms were recorded by absorbance at 280 nm. Individual AH fusion proteins and equimolar mixtures of AH-ST and AH-SC were analyzed. Before SEC, protein samples were mixed with SDS to a final concentration of 0.5% (w/v). Where indicated, samples were additionally heated at 95 °C for 8 min prior to analysis. A sample volume of 10 μL was used for each SEC analysis.

### Cryo-EM images and structural analysis

A 3 μL aliquot of the sample solution was applied onto a glow-discharged custom-made ultra-thin carbon coated Quantifoil R1.2/1.3 200 mesh Cu grids (Quantifoil, Jena, Germany). Then, the grid was blotted and rapidly vitrified in liquid ethane using a Vitrobot Mark IV (Thermo Fisher Scientific, Waltham, MA, USA).

Cryo-EM data were acquired automatically using SerialEM program on a CRYO ARM 300 II transmission electron microscope (JEOL, Tokyo, Japan), operated at 300 kV accelerating voltage. Images were obtained by the CDS mode of the K3 camera direct electron detector (Gatan, Pleasanton, USA). The total exposure dose on the specimen was 40 e^−^Å^−2^. The details of the data acquisition condition are described in Table S3.

The obtained movie frames were motion corrected with the program Patch Motion Correction implemented in cryoSPARC3.4, with dose weighting. CTFs of the motion-corrected micrographs were also estimated with the program Patch CTF Estimation implemented in cryoSPARC3.4. A total of 7196 and 3945 micrographs were subjected to subsequent image processing for WT and K237C AH-ST/SC respectively. Particle picking was performed with the program crYOLO1.9.3. Further image processing was performed with cryoSPARC. A total of 385,089 and 811,025 particles were extracted and subjected to 2D classification for WT and K237C AH-ST/SC respectively. Particles from 2D classes, which converged as AH-dimers (349,752 and 579,248 particles for WT and K237C AH-ST/SC) were refined, which resulted in final maps with a resolution of 2.20 Å for both WT and K237C AH-ST/SC. The detailed image processing conditions are shown in Figure S4 and S5.

Atomic coordinates were constructed manually with the program COOT using crystal structure of AH (PDB ID 3ANS) as a starting model. Structure refinement was perfomed using phenix.real_space_refine. Structural superposition and analysis were performed with the program UCSF ChimeraX 1.12. The two structures were superimposed on their lower heptamers with the MatchMaker tool (r.m.s.d. 0.211 Å), and the relative rotation of the upper heptamer was determined by separately superimposing the upper heptamers and measuring the resulting transformation. Interface distances were measured between the modelled side-chain atoms of the two opposing chains at the heptamer–heptamer interface. Atomic coordinates and cryo-EM maps were deposited in the PDB (8JX2, 8JX3) and EMDB (EMD-36688, EMD-36689) for WT and K237C AH-ST/SC respectively. Statistics of structure refinement are summarized in Table S4.

### Preparation and quantification of GUVs

GUVs were prepared using the phase-transfer method^40^. Lipid mixtures with the desired compositions were dissolved in 500 μL liquid paraffin and mixed with 20 μL inner aqueous solution to generate water-in-oil emulsions. For calcein leakage assay, lipid mixtures were composed of POPC 44 (mol%), cholesterol (55.5 mol%) and DSPE-PEG2k (0.5 mol%). When POPG was added, lipid mixtures were changed to POPC (85.5 mol%), POPG (9.3 mol%), cholesterol (4.7 mol%) and DSPE-PEG2k (0.5 mol%) instead. For confocal microscopy, DSPE-PEG2k was replaced with biotin-DSPE-PEG2k to immobilize GUVs on the cover glass coated with streptavidin. Unless otherwise specified, the inner buffer consisted of 20 mM HEPES-KOH (pH 7.6), 250 mM potassium glutamate, 18 mM magnesium acetate, 120 mM glucose, and 280 mM sucrose. The outer buffer consisted of 20 mM HEPES-KOH (pH 7.6), 250 mM potassium glutamate, 18 mM magnesium acetate, and 400 mM glucose. The water-in-oil emulsion was gently layered onto 300 μL of outer buffer in a 1.5 mL microcentrifuge tube and centrifuged at 10,000 × g for 5 min at 25 °C. Following centrifugation, the upper oil phase was carefully removed, and the resulting GUV pellet was washed three times with outer buffer before being resuspended in 50 μL of outer buffer.

For specific experiments, fluorescent reporters (calcein, OA647, SNAP-GFP, mCherry, Cascade Blue conjugated-dextran or caged-EtBr) or nucleic acid cargo (λDNA) were encapsulated in the inner solution as indicated. The composition of the inner and outer buffers was otherwise maintained unchanged.

The lipid concentration of purified GUV suspensions was quantified using a Phospholipid Assay Kit (Sigma-Aldrich). Prior to use, GUV suspensions were diluted with outer buffer to a final lipid concentration of 500 μM.

### FCM analysis

FCM measurements were performed using a FACS Aria III flow cytometer (BD Biosciences, San Jose, CA, USA). Samples were diluted with outer buffer immediately prior to measurement. A total of 10,000 events were collected for each sample. Data for calcein leakage assay were obtained using the following setting: calcein was measured using a 488-nm blue laser and a 530/30 filter (429 V); AF647 was measured using a red 633-nm laser and a 660/20 filter (429 V); FS and SS was measured with a voltage of 3 and 380, respectively. Thresholds were set at a value of 500 for both FSC and SSC, and 1000 for AF647. Data for vesicle linkage assay were obtained using the following setting: SNAP-GFP was measured using a 488-nm blue laser and a 530/30 filter (544V); AF647 was measured using a red 633-nm laser and a 660/20 filter (582); FS and SS was measured with a voltage of 5 and 402, respectively. Thresholds were set at a value of 500 for both FSC and SSC. Data analysis was performed using FlowJo software (v10.9.0, BD Life Sciences).

### Confocal fluorescence microscopy

Confocal fluorescence images were acquired using a Zeiss laser-scanning confocal microscope (Carl Zeiss Microscopy GmbH, Jena, Germany). Prior to observation, coverslips were first coated with 0.1 mg/ml neutravidin for 1hr, followed by blocking with 1% (w/v) BSA for 1hr. GUV samples were mounted between two cover glasses and sealed with liquid paraffin to minimize sample drift during observation.

### Calcein leakage assay

Calcein leakage from GUVs was evaluated by FCM. GUVs encapsulating 10 μM calcein (FUJIFILM Wako Pure Chemical Corporation, Osaka, Japan) and 5 μM OA647 (Thermo Fisher Scientific, Waltham, MA, USA) were incubated with AH fusion proteins at final protein concentrations ranging from 0 to 4 μM. The final lipid concentration was 250 μM.

Samples were incubated at 25 °C for 0.5 - 24 h. Calcein retention was quantified as the geometric mean of the calcein/OA647 fluorescence; a decrease in this value indicates the decrease in calcein concentration.

### Droplet interface bilayer measurements

Single-channel currents through AH nanopores were measured using a droplet-interface-bilayer (DIB) microdevice consisting of two 2.0-mm-diameter chambers separated by a parylene C film (Figure S8). Ag/AgCl electrodes positioned in the two chambers were connected to a Pico patch-clamp amplifier (Tecella, Foothill Ranch, CA, USA). A planar lipid bilayer was first prepared using the droplet contact method within the microdevice^41^. Two lipid monolayers contacted each other and formed a bilayer through a 100 µm-diameter hole in a 5 µm-thick parylene C (polychloro-p-xylylene, Parylene Japan, Japan) film. Each chamber received 0.3 µL of 20 mg/mL DPhPC dissolved in *n-*decane (FUJIFILM Wako Pure Chemical Industries, Ltd. (Wako), Osaka, Japan) into each chamber. A volume of 4.7 µL of the solution containing the AH nanopore, 1 M KCl, 10 mM MOPS was added to the ground chamber, and 4.7 µL of the solution containing 1 M KCl, 10 mM MOPS was added to the recording chamber. The channel current was detected with an 8 kHz low-pass Bessel filter and a sampling rate of 40 kHz. A constant voltage of +100 mV was applied to the recording chamber. Currents were low-pass filtered at 8 kHz using a Bessel filter and sampled at 40 kHz. Current traces were analyzed using Clampfit v11.3 (Molecular Devices). Single-channel conductance was calculated as G = ΔI/V, where ΔI is the current-step amplitude and V is the applied voltage.

### Vesicle linkage assay

Two GUV populations were prepared, with one population encapsulating 5 μM SNAP-GFP and the other encapsulating 5 μM OA647 (or mCherry for confocal microscopy). AH-ST- or AH-SC-functionalized GUVs were generated by incubating the corresponding AH fusion proteins with GUVs at final protein concentrations of 1 μM for 2 h at 25 °C to allow membrane insertion.

Equal volumes of the two independently functionalized GUV populations were mixed and incubated for an additional 3 h at 25 °C. For the ST/ST and SC/SC control groups, both GUV populations were functionalized with the same AH fusion protein. For the ST/SC group, one GUV population was functionalized with AH-ST and the other with AH-SC. Samples were subsequently diluted with outer buffer and analyzed by FCM. The fraction of double-positive (Q2) events (Figure 4a) was used as an operational measure of association between the two fluorescently distinct GUV populations.

### Synthesis and characterization of caged-EtBr

To a mixture of a DMSO solution of PC Alkyne-PEG4-NHS (100 mM, 15 μL), an aqueous solution of EtBr (25.4 mM, 147.6 μL), and DMSO (62.4 μL), an aqueous solution of pyridine-HCl buffer (1 M, pH 5.0, 225 μL) was added. After shaking at 37 °C for 3.5 h, the reaction was quenched by the addition of 0.38% formic acid solution and MeOH. The crude mixture was purified by reverse-phase HPLC (XBridge™ C18, 5 μm, 10 × 50 mm; flow rate, 3.0 mL min^-1^ with a linear gradient of 0-100% MeOH in 0.38% formic acid over 15 min) to afford alkyne-containing caged EtBr (822.1 nmol, 55% yield). The product was identified by MALDI-TOF-MS (calcd for [M]^+^, m/z 852.4; found, m/z 852.4).

The alkyne-containing caged EtBr was then conjugated with mPEG_5000_-azide by copper(I)-catalyzed azide-alkyne cycloaddition (CuAAC). To a mixture of a DMSO solution of alkyne-containing caged EtBr (2 mM, 200 μL) and an aqueous solution of mPEG_5000_-azide (10 mM, 80 μL), aqueous solutions of THPTA (50 mM, 80 μL), sodium ascorbate (200 mM, 200 μL), and CuSO_4_ (10 mM, 80 μL) were added. After incubation with shaking at 30 °C for 1 h, the reaction was quenched by the addition of 0.38% formic acid solution. The crude mixture was purified by reverse-phase HPLC (XBridge™ C18, 5 μm, 10 × 50 mm; flow rate, 3.0 mL min^-1^ with a linear gradient of 0-100% MeOH in 0.38% formic acid over 15 min) to afford mPEG_5000_-conjugated caged EtBr (323.0 nmol, 81% yield). The product was characterized by MALDI-TOF-MS. Because mPEG_5000_-azide was a polydisperse PEG reagent, the observed product peak was assigned to the mPEG_5000_-conjugated caged EtBr species containing 108 ethylene glycol units (calcd for [M]^+^, m/z 5708.3; found, m/z 5708.3).

### Photo-controlled cargo release and uptake assay

Two GUV populations were prepared for the chemical transport assay. Sender GUVs encapsulated 5 μM OA647 and 100 μM caged-EtBr, whereas receiver GUVs encapsulated 1mg/ml Dextran, Cascade Blue^TM^ (10,000 MW, Anionic, Lysine Fixable Thermo Fisher Scientific, Waltham, MA, USA) and λDNA (Takara Bio Inc., Kusatsu, Shiga, Japan) at a base-pair concentration of 100 μM. AH-ST- and AH-SC-functionalized GUVs were prepared as described for the vesicle-linkage assay. Following vesicle linkage, the entire samples were irradiated with 365-nm UV light (LED 365 nm, 3 cm between LED and the samples; power 25%, 84.9 mW/cm^-2^) for 3 min to induce uncaging of EtBr from caged-EtBr.

Receiver GUVs were identified by cascade blue fluorescence and imaged by confocal fluorescence microscopy before and after UV irradiation. EtBr fluorescence was quantified for individual receiver GUVs. The fluorescent increase for each receiver GUV was calculated as Δ*F* = *F*_after_ − *F*_before_. For analysis of the ST/ST controls, receiver GUVs were additionally classified according to their apparent spatial relationship with sender GUVs. A receiver GUV was classified as adjacent when its membrane appeared to be in direct contact with an OA647-positive sender GUV in the same focal plane, as judged from the OA647 and cascade blue channels. All other receiver GUVs were classified as non-adjacent.

### Data analysis

Flow cytometry data were analyzed with FlowJo (FlowJo, LLC, Ashland, OR, USA). Flow-cytometry fluorescence measurements were summarized using the geometric mean or median of the gated event population as specified for each experiment. Confocal fluorescence measurements were quantified at the individual-GUV level as described above. Statistical analysis was performed using OriginPro 2026b. All relevant information on statistical analysis, including *p*-values, the number of GUVs that were analyzed and the statistical test used can be found in the respective figure legends. No statistical method was used to predetermine sample size. No data were excluded from the analyses and experiments were not randomized. The investigators were not blinded to allocation during experiments and outcome assessment.

## Acknowledgement

We thank Hiroki Ijima from the Graduate School of Life Sciences, Tohoku University for assistance with cryo-EM data processing and for preparing Figures S4 and S5.

## Grants

This study was supported by Human Frontier Science Program Grant Number RGP003/2023 (T.M.), JSPS KAKENHI Grant Numbers 22K21344 (T.M.), 21H05228 (T.M.) and JST ASPIRE Grant Number JPMJAP24B4 (T.M.).

## Notes

### Competing Interest Statement

The authors have declared no competing interest.

