## Supplementary Information for "Selective Recognition, Covalent Linkage and Chemical Exchange of Artificial Cells through Engineered α-Hemolysin"

\*Corresponding authors

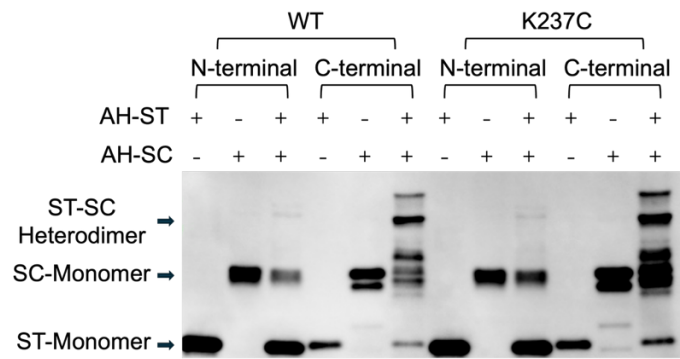

**Figure S1. Covalent ST-SC complex formation requires C-terminal fusion to AH.** Western blot analysis of AH variants carrying ST or SC at either the N- or C- terminus. Proteins were synthesized individually using the PURE system, mixed pairwise, and detected with an anti-His6 antibody-HRP conjugate. A band corresponding to the covalent ST-SC heterodimer was detected only for the C-terminal fusions, in both the WT and K237C backgrounds.

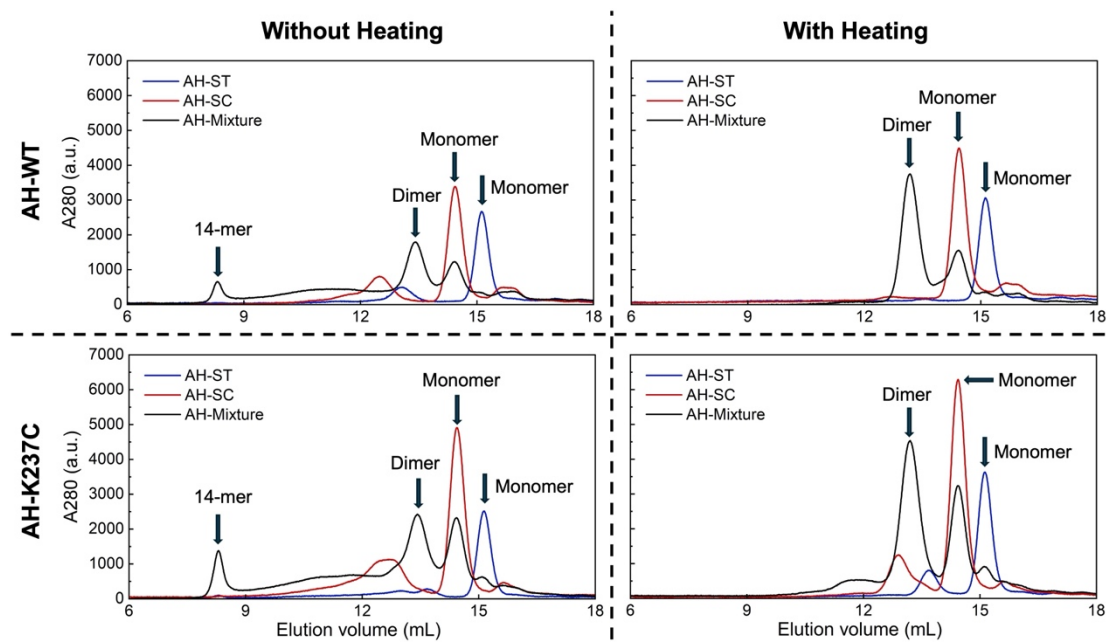

**Figure S2. Size-exclusion chromatography of AH fusion proteins and equimolar mixtures of ST and SC proteins.** Both WT and K237C constructs, AH-ST alone (blue), AH-SC alone (red), and equimolar mixtures of the two (black) were analyzed on a Superose 6 Increase 10/300 GL column in the presence of 0.5% (w/v) SDS. Samples were analyzed either without heating or after heating at 95°C for 8min before the analysis. Absorbance at A280 nm is plotted against elution volume. An early-eluting peak between 6-9 mL was observed only in the non-heated mixtures and was assigned to the 14-mer assembly; this peak disappeared upon heating, whereas the peak assigned to the covalent ST-SC heterodimer between 12-15 mL was retained.

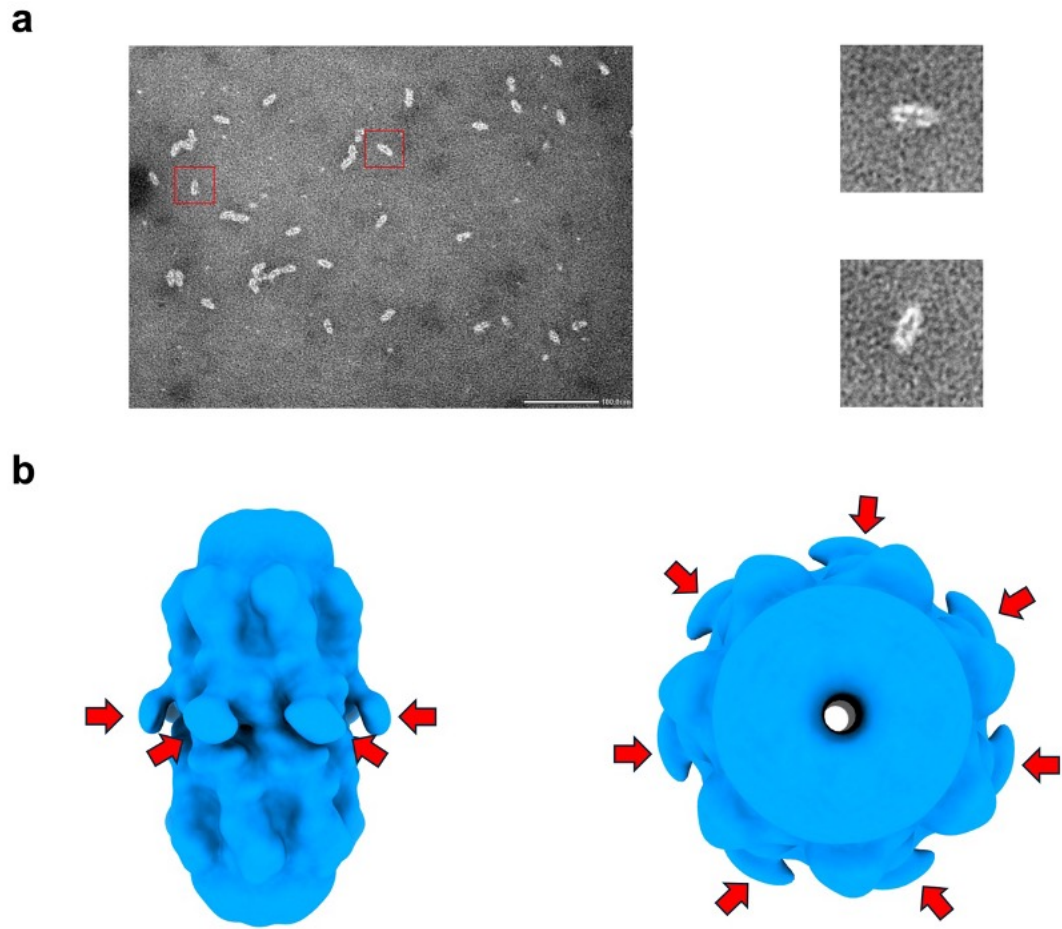

**Figure S3. Particle images and unassigned densities at the heptamer–heptamer interface.** **a)** Representative negative stain TEM (JEM1400-Plus, JEOL) of the WT AH-ST/AH-SC mixture. Elongated, dumbbell-shaped particles consistent with head-to-head 14-mer assemblies were abundant. Two particles are boxed in red and shown enlarged on the right. Scale bar: 100 nm. **b)** Gaussian-filtered cryo-EM map of the WT 14-mer assembly, shown in side view (left) and top view (right). Weak, unresolvable densities were observed at seven equivalent positions around the interface between the two opposing heptamers (red arrows), consistent with the presence of flexibly tethered ST and SC domains that could not be modelled.

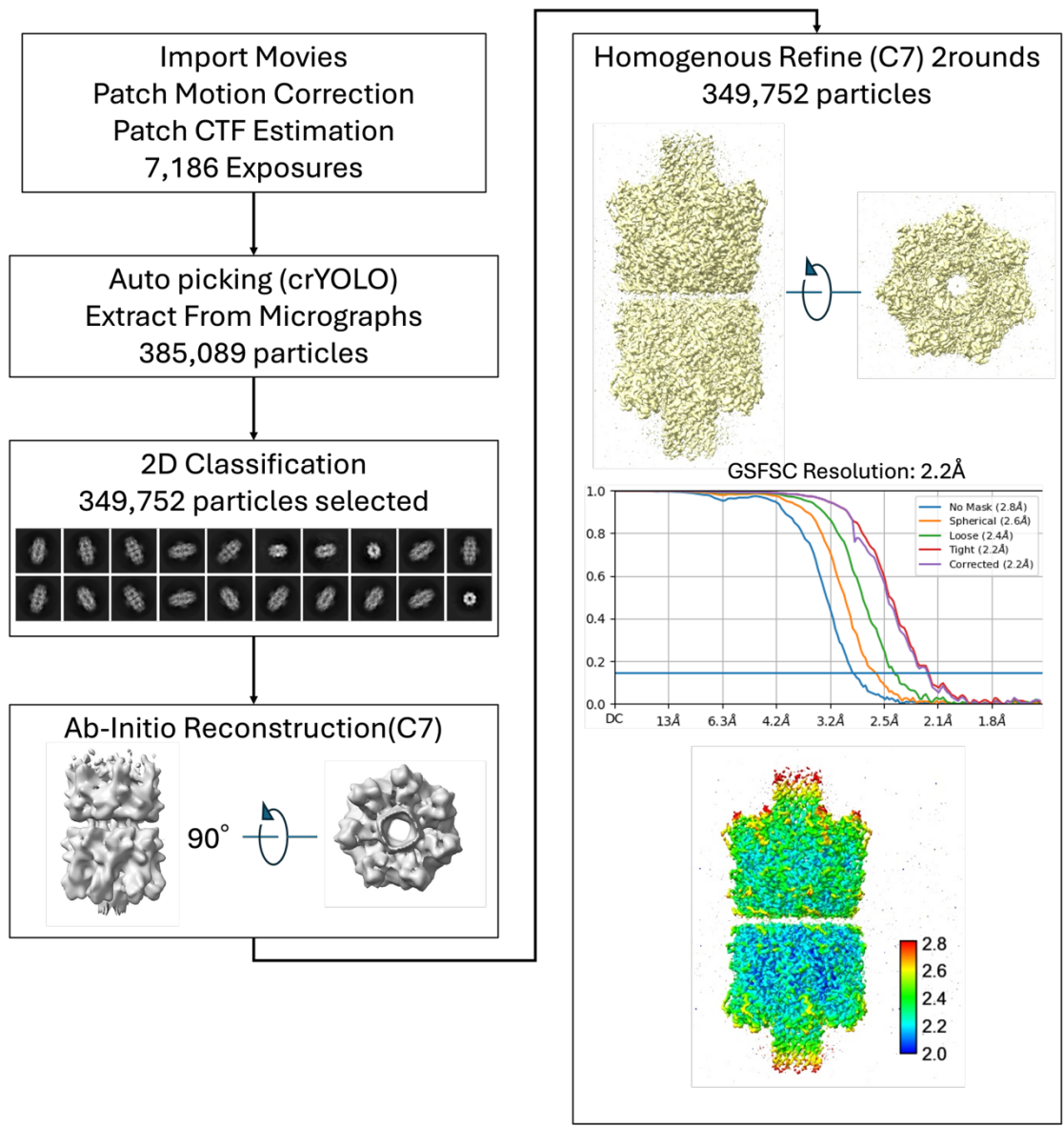

**Figure S4. Cryo-EM data processing pipeline of AH-WT.** Movies were imported and subjected to patch motion correction and patch CTF estimation in cryoSPARC. Particles were auto-picked with crYOLO and extracted (385,089 particles), and 349,752 particles were selected after 2D classification. The 3D map was generated with C7 symmetry imposed, followed by two rounds of homogeneous refinement with C7 symmetry. The gold-standard FSC curves (GSFSC) are shown, giving a corrected resolution of 2.2 Å at the FSC = 0.143 criterion, together with the local resolution estimate colored from 2.0 to 2.8 Å.

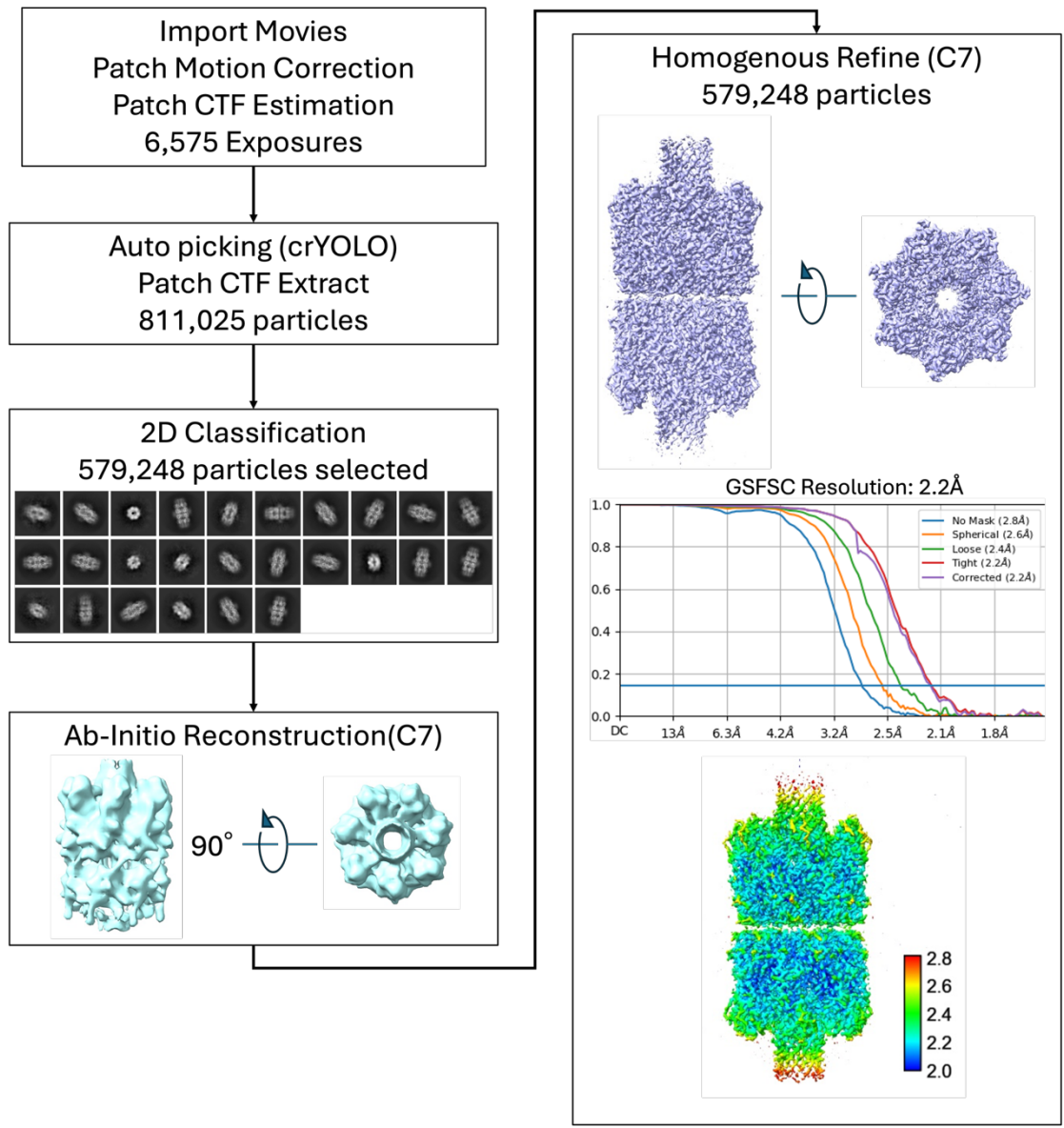

**Figure S5. Cryo-EM data processing pipeline of AH-K237C.** Movies were imported and subjected to patch motion correction and patch CTF estimation in cryoSPARC. Particles were auto-picked with crYOLO and extracted (811,025 particles), and 579,248 particles were selected after 2D classification. The 3D map was generated with C7 symmetry imposed, followed by two rounds of homogeneous refinement with C7 symmetry. The gold-standard FSC curves (GSFSC) are shown, giving a corrected resolution of 2.2 Å at the FSC = 0.143 criterion, together with the local resolution estimate colored from 2.0 to 2.8 Å.

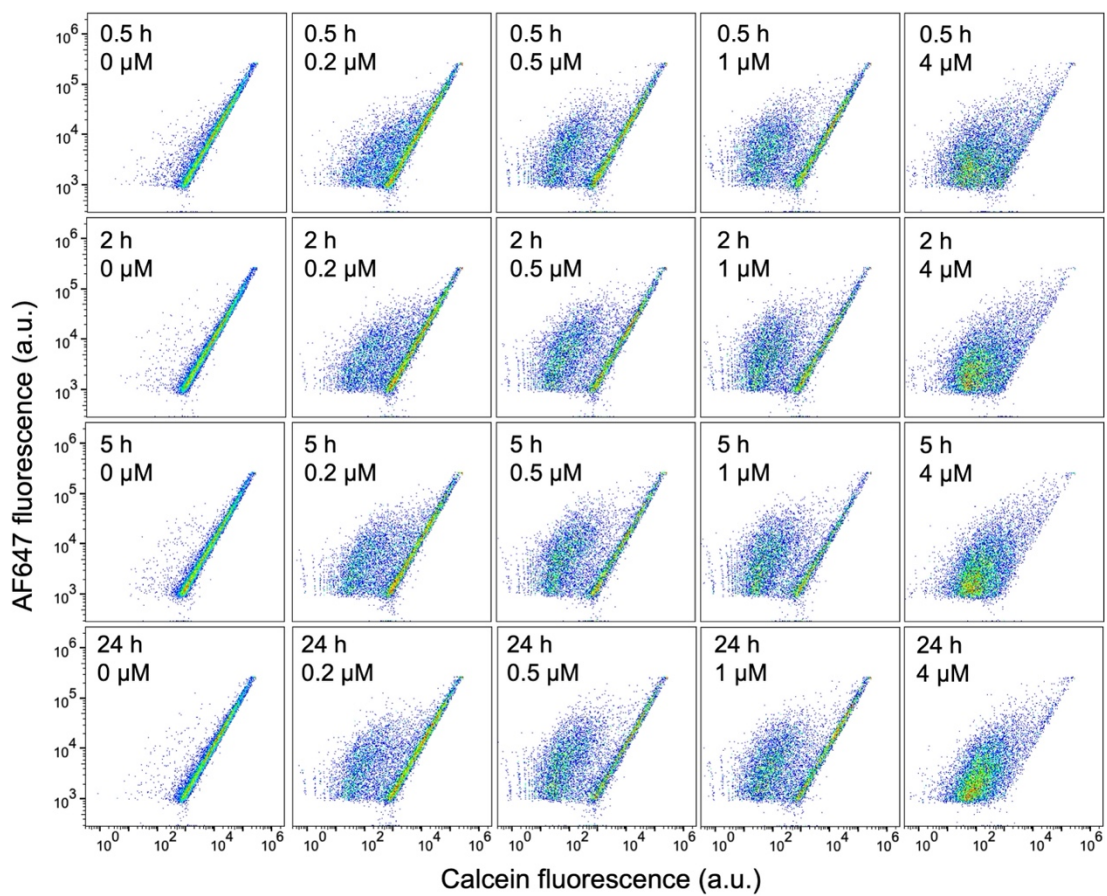

**Figure S6. Effects of AH concentration and incubation time on calcein retention in GUVs.** Two-dimensional FCM plots of calcein versus AF647 (OA647) fluorescence for GUVs incubated with WT-ST at 0, 0.2, 0.5, 1, and 4 $\mu\text{M}$ for 0.5, 2, 5 and 24 h at 25 $^{\circ}\text{C}$. OA647 was retained within the GUV lumen and served as a per-vesicle internal reference, so pore-mediated calcein release appeared as a shift along the calcein axis with comparatively stable OA647 fluorescence. Increasing protein concentration produced a progressive shift, whereas prolonging incubation beyond 5 h produced little further change. WT-ST is shown as a representative example; the same trend was observed for all four AH proteins and is quantified in Figure S7. Each plot represents 10,000 events from one representative preparation. Note that AH treated GUVs show binary distribution, where one has low calcein concentration and the other having the initial calcein concentrations.

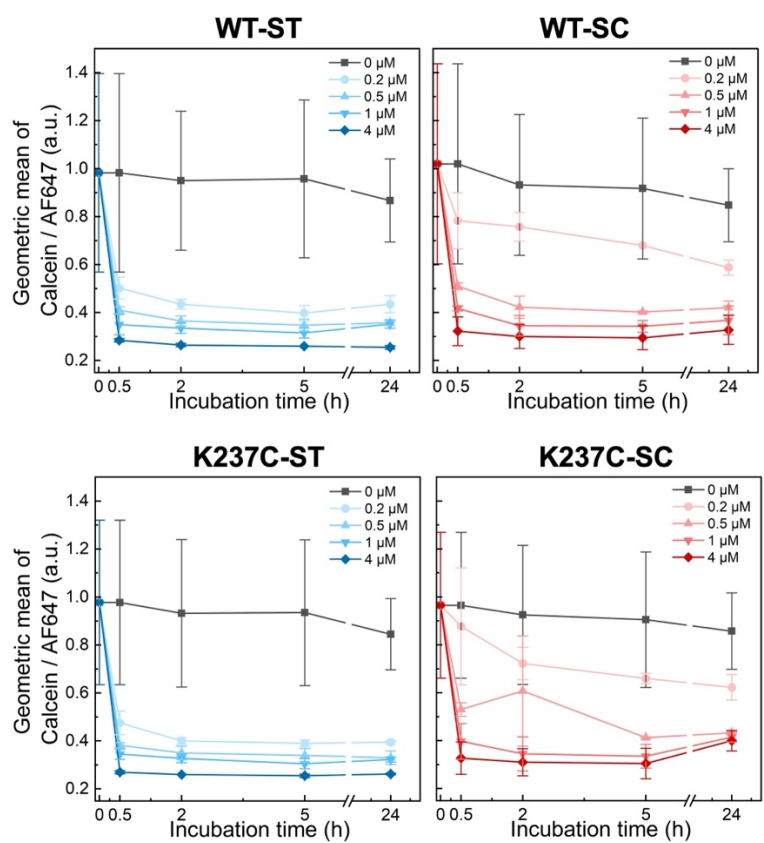

**Figure S7. Time-dependent calcein leakage induced by engineered AH fusion proteins.** GUVs were incubated with WT-ST, WT-SC, K237C-ST, or K237C-SC at 0, 0.2, 0.5, 1, and 4 $\mu\text{M}$ at 25 $^{\circ}\text{C}$, and calcein leakage was measured by FCM at 0, 0.5, 2, 5 h, and 24hr, respectively. The geometric mean of the calcein/AF647 fluorescence ratio for individual events was plotted against incubation time. Data of 0 $\mu\text{M}$ AH was used as the time 0 data. Data are mean ± SD from two independent GUV preparations, each measured in duplicate. For all four AH proteins, the ratio approached a plateau after approximately 5 h, which was therefore adopted for all subsequent experiments.

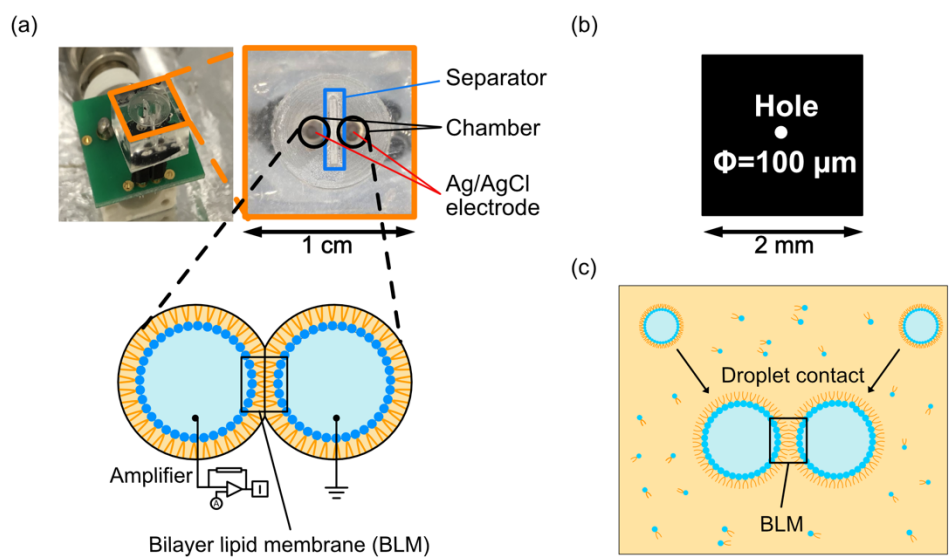

**Figure S8. Microdevice for channel current measurement.** Measurements were done essentially as described previously<sup>1</sup>. **a)** Photograph of the device (left) and enlarged view of the measurement region (right), consisting of two 2.0 mm-diameter chambers separated by a separator, each equipped with an Ag/AgCl electrode connected to a patch-clamp amplifier. The schematic below illustrates the droplet contact method, in which two lipid monolayers meet to form a bilayer lipid membrane (BLM) at the interface. **b)** The separator is a 5 $\mu\text{m}$-thick parylene C film containing a 100 $\mu\text{m}$-diameter hole through which the bilayer is formed. **c)** Schematic of bilayer formation: lipid monolayers self-assemble at the interface between the aqueous droplets and the surrounding n-decane, and a BLM forms where the two droplets come into contact.

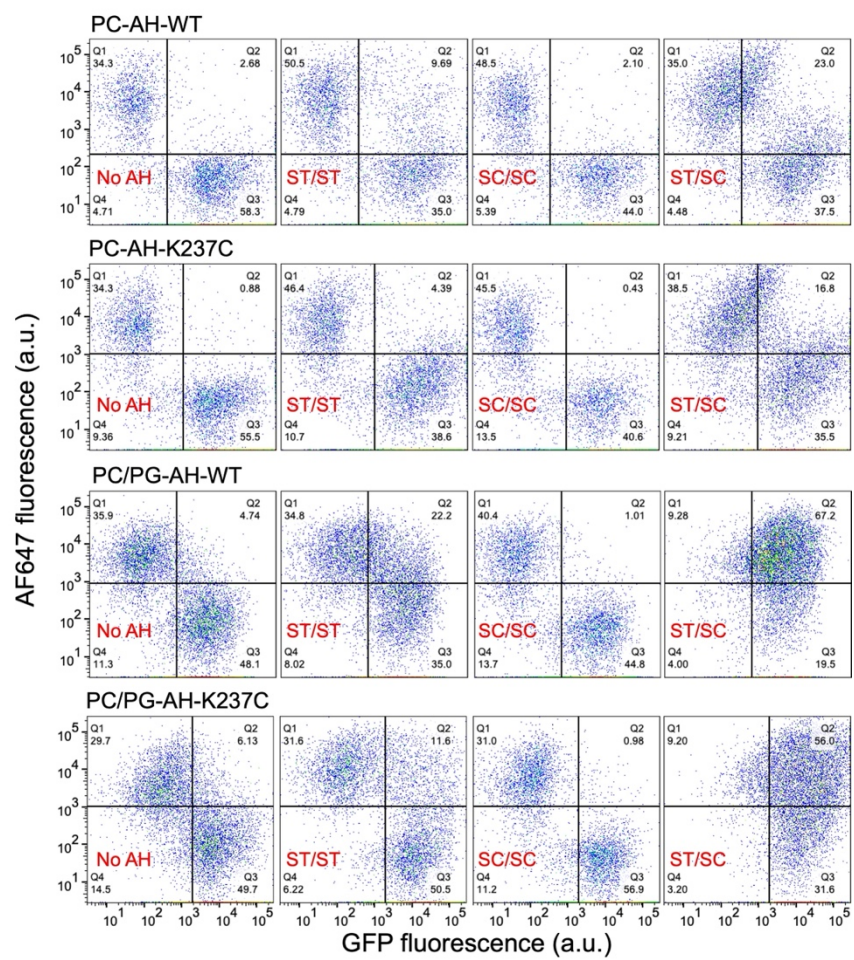

**Figure S9. Effect of membrane lipid composition on ST-SC-mediated GUV linkage.** Representative FCM plots corresponding to the quantitative data shown in Figure 4b. Two-dimensional FCM plots of GFP versus AF647 fluorescence for GUVs prepared with PC or PC/PG and functionalized with the indicated AH proteins. Quadrant gates were set independently for each replicate using the no-AH control. Q2 contains double-positive events containing both GFP and AF647 fluorescence. Incorporation of PG increased the Q2 fraction for the ST/ST and ST/SC conditions, while the SC/SC control remained unchanged; the ST/SC group consistently exceeded both controls. Each plot represents 10,000 events from one representative preparation.

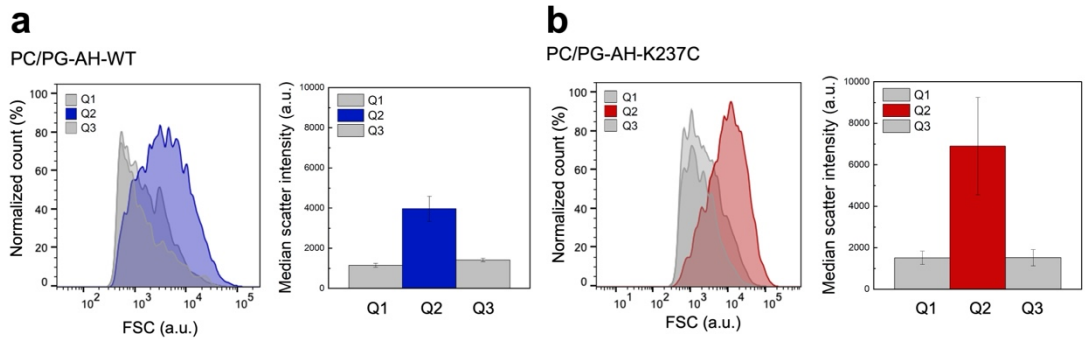

**Figure S10. Forward scatter of gated populations.** Histogram and median scattering intensities of FSC for the Q1, Q2 and Q3 populations of the ST/SC sample in **a)** PC/PG-AH-WT and **b)** PC/PG-AH-K237C. Populations containing fewer than 200 events were excluded from this analysis; the SC/SC double-positive population fell below this threshold and is therefore not shown. Histogram of FSC is from one representative preparation. Bars show the mean ± SD of the median FSC values obtained from two WT and three K237C independent GUV preparations.

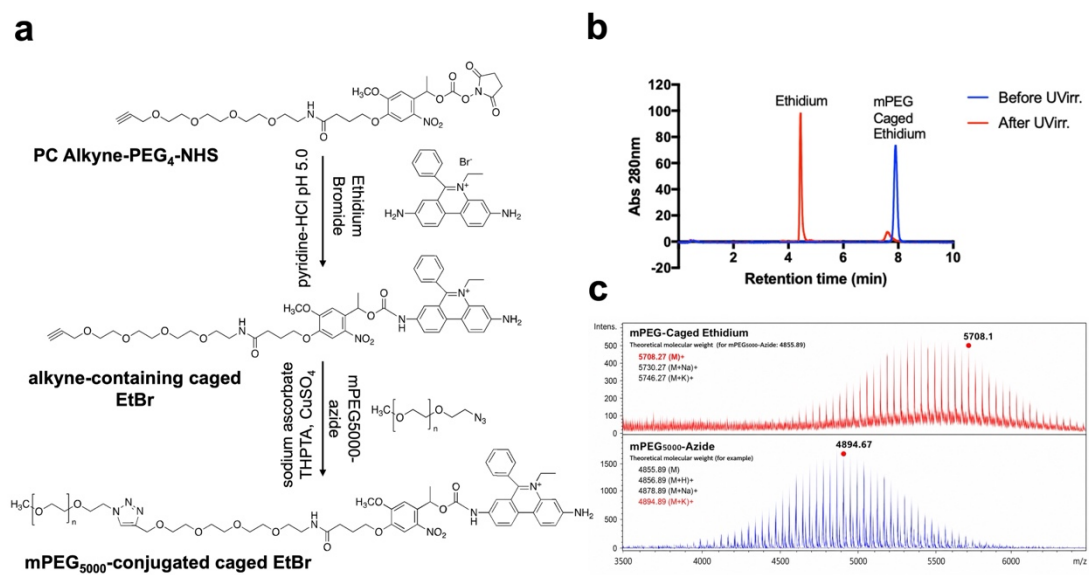

**Figure S11. Synthesis and characterization of mPEG<sub>5000</sub>-conjugated caged EtBr.**

**a)** Synthetic scheme of mPEG<sub>5000</sub>-conjugated caged EtBr. **b)** Overlaid reversed-phase HPLC chromatograms of caged EtBr before UV-irradiation (blue) and after UV-irradiation with 365 nm UV light (2.96 mW/cm<sup>2</sup>) for 10 min (red). The main red peak was assigned to the photolysis product, ethidium. **c)** MALDI-TOF-MS analysis of mPEG<sub>5000</sub>-conjugated caged EtBr (upper plot). Since mPEG<sub>5000</sub>-azide was a polydisperse PEG reagent (lower plot), one of the observed product peaks was assigned as a representative based on the corresponding mPEG<sub>5000</sub>-azide species observed at $m/z$ 4894.9 [M+K]<sup>+</sup>, which contained 108 ethylene glycol units. The calculated mass of the corresponding caged EtBr species with the aforementioned PEG was $m/z$ 5708.3 [M]<sup>+</sup>, consistent with the observed peak.

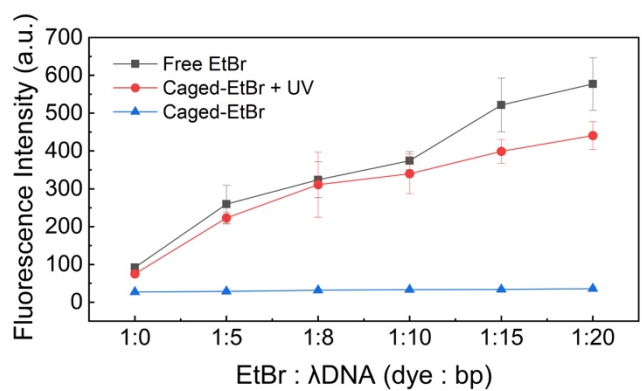

**Figure S12. Bulk validation of UV-triggered EtBr release and DNA binding.** Fluorescence intensity changes of free EtBr and caged EtBr before and after UV irradiation measured in a 384-well plate. EtBr concentration was fixed to 2 μM, and the concentration of λDNA (base-pair concentration) was adjusted to 0, 10, 16, 20, 30 and 40 μM respectively according to the ratio. Fluorescence was recorded at $\lambda_{\text{ex}}$ = 510 nm and $\lambda_{\text{em}}$ = 610 nm. A pronounced fluorescence increase was observed only when caged EtBr and λDNA were both present and the sample was UV irradiated, confirming that photo-uncaging is required for λDNA binding. Data are mean ± SD from three independent experiments, each measured in triplicate (n = 3).

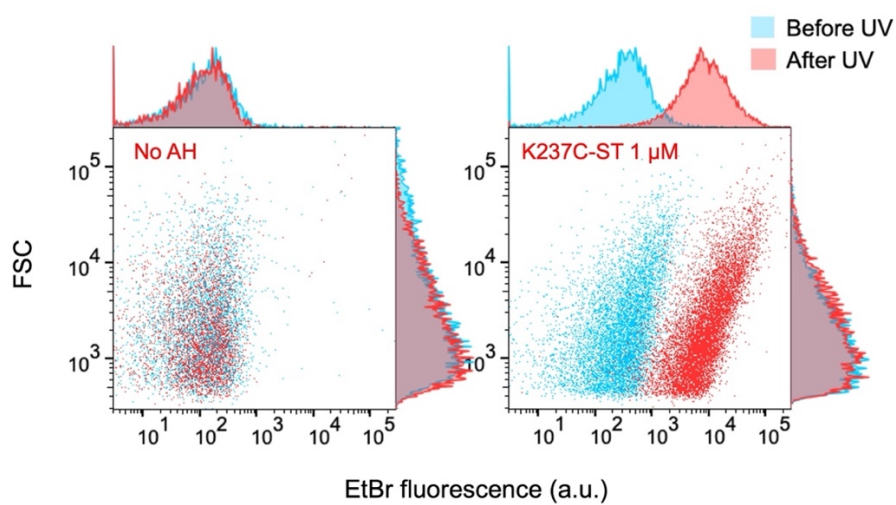

**Figure S13. AH-dependent accumulation of photo-released EtBr in λDNA-containing GUVs.** GUVs encapsulating λDNA were incubated either without AH (left) or with the 1 μM K237C-ST for 2 h at 25 $^{\circ}$C to allow membrane insertion. Caged-EtBr (100 μM) was then added to the external solution of GUVs, and the samples were analyzed by FCM before and after UV irradiation. EtBr fluorescence shifted to higher intensity in the presence of AH and after UV irradiation, indicating that photo-uncaging released free EtBr from the PEG-conjugated caged precursor entered the GUV lumen through AH nanopores and bound the encapsulated λDNA. A modestly higher pre-irradiation EtBr signal was also observed in the K237C-ST sample and may reflect trace amounts of free EtBr present in the caged-EtBr preparation that entered the GUVs through AH pores. Each plot represents 10,000 events from a single experiment.

**Table S1. Primers used for AH construction in this study.**

| **Primer** | **Sequence (5' → '3)** |
| --- | --- |
| T7F | TAATACGACTCACTATAGGG |
| T7R | ATGCTAGTTATTGCTCAGCGG |
| K237C-FP | ATTACTATGGATAGATGCGCATCCAAACAACAAAC |
| K237C-RP | GTTTGTTGTTTGGATGCGCATCTATCCATAGTAAT |
| AH-NF | GCAGATTCTGATATTAATATTAAAACCGG |
| AH-NR | CATGGTATATCTCCTTCTTAAAGTTAAAC |
| AH-CF | GGGGGTTCTCATCATCATC |
| AH-CR | ATTTGTCATTTCTTCTTTTTCCCAATC |
| AHST-NF | TATCGTGATGGTGGATGCATACAAACGCTATAAAGGATCCAGT<br>GGTAGCGCAGATTCTGATATTAATATTAAAACCGG |
| AHST-NR | CATCCACCATCACGATATGCGGAACACCACGCATGGTATATC<br>TCCTTCTTAAAGTTAAAC |
| AHST-CF | GTTCCGCATATCGTGATGGTGGATGCATACAAACGCTATAAA<br>GGGGGTTCTCATCATCATC |
| AHST-CR | CCATCACGATATGCGGAACACCACGGCTACCACTGGATCCAT<br>TTGTCATTTCTTCTTTTTCC |
| SC3-GS-F | AGAAGGAGATATACCATGTCGTACTACCATCACCATCACC |
| SC3-GS-R | ATATTAATATCAGAATCTGCGCTACCACTGGATCCAGTATGAG<br>CGTCACCTTCAGTTG |
| GS-SC3-F | GAAGAAATGACAAATGGATCCAGTGGTAGCTCGTACTACCAT<br>CACCATCACC |
| GS-SC3-R | GAGCGGATCCACAATTTAAGTATGAGCGTCACCTTCAGTTG |
| pET28b-F | TGAGATCCGGCTGCTAACAAAG |
| pET28b-R | GGTATATCTCCTTCTTAAAGTTAAACAAAATTA |
| 2fix-to-<br>pET28b-F | TAAGAAGGAGATATACCATGGCAGATTCTGATATTAATATT |
| 2fix-to-<br>pET28b-R | GTTAGCAGCCGGATCTCATTAATGATGATGATGATGATGAG |

**Table S2. Plasmids used in this study**

| Plasmid | Description | Selection marker |
| --- | --- | --- |
| pIVEX2.3d-AH-6CHis-2fix | α-Hemolysin(G122S/K147R)-6CHis | AmpR |
| pIVEX2.3d-AH-6CHis-2fix-K237C | AH-G122S/K147R/K237C |  |
| pIVEX2.3d-AH-6CHis-2fix-N-ST | AH-G122S/K147R-N-terminal SpyTag003 |  |
| pIVEX2.3d-AH-6CHis-2fix-N-SC | AH-G122S/K147R-N-terminal SpyCatcher003 |  |
| pIVEX2.3d-AH-6CHis-2fix-C-ST | AH-G122S/K147R-C-terminal SpyTag003 |  |
| pIVEX2.3d-AH-6CHis-2fix-C-SC | AH-G122S/K147R-C-terminal SpyCatcher003 |  |
| pIVEX2.3d-AH-6CHis-2fix-K237C-N-ST | AH-G122S/K147R/K237C-N-terminal SpyTag003 |  |
| pIVEX2.3d-AH-6CHis-2fix-K237C-N-SC | AH-G122S/K147R/K237C-N-terminal SpyCatcher003 |  |
| pIVEX2.3d-AH-6CHis-2fix-K237C-C-ST | AH-G122S/K147R/K237C-C-terminal SpyTag003 |  |
| pIVEX2.3d-AH-6CHis-2fix-K237C-C-SC | AH-G122S/K147R/K237C-C-terminal SpyCatcher003 |  |
| pET28b-AH-6CHis-2fix-C-ST | AH-G122S/K147R-C-terminal SpyTag003 | KanR |
| pET28b-AH-6CHis-2fix-C-SC | AH-G122S/K147R-C-terminal SpyCatcher003 |  |
| pET28b-AH-6CHis-2fix-K237C-C-ST | AH-G122S/K147R/K237C-C-terminal SpyTag003 |  |
| pET28b-AH-6CHis-2fix-K237C-C-SC | AH-G122S/K147R/K237C-C-terminal SpyCatcher003 |  |
| pET28b | Cloning vector |  |

**Table S3. Cryo-EM Data collection for AH-WT and AH-K237C**

| **Data collection** | **AH-WT** | **AH-K237C** |
| --- | --- | --- |
| Microscope | CRYO ARM 300 II | CRYO ARM 300 II |
| Voltage (kV) | 300 | 300 |
| Detector | GATAN K3 (6k x 4k) | GATAN K3 (6k x 4k) |
| Electron exposure (e<sup>-</sup>/Å<sup>2</sup>) | 40 | 40 |
| Magnification (nominal) | 60,000 | 60,000 |
| Pixel size (Å/pix) | 0.788 | 0.788 |
| Defocus range (μm) | 1.5 to 2.5 | 1.5 to 2.4 |
| Number of movies/micrographs | 7196 | 3945 |
| Map dimensions | 400, 400, 400 | 400, 400, 400 |
| Symmetry imposed | C7 | C7 |
| Number of particles | 349,752 | 579,248 |
| FSC threshold | 0.143 | 0.143 |

**Table S4. Cryo-EM Data refinement for AH-WT and AH-K237C**

| **Refinement** | **AH-WT** | **AH-K237C** |
| --- | --- | --- |
| Initial model (PDB) | 3ANS | 3ANS |
| Non-hydrogen atoms | 32,872 | 32,826 |
| Protein residues (modeled) | 4,102(14×293) | 4,102(14×293) |
| Ligands | None | None |
| Clashscore | 4 | 4 |
| Poor rotamers (%) | 3.3(123/3640) | 2.1 (70/3347) |
| Ramachandran favored (%) | 98 | 98 |
| Ramachandran allowed (%) | 2 | 2 |
| Ramachandran outliers (%) | 0 | 0 |
| Bond length RMSZ | 0.12 | 0.28 |
| Bond angle RMSZ | 0.30 | 0.36 |
| Map-model Q-score (all) | 0.6740 | 0.681 |
| Atom inclusion (backbone / all) | 99% / 94% | 99% / 95% |

#### Note S1. Theoretical number of AH heptamer bound to each GUV

The theoretical number of membrane-inserted AH heptamers per GUV was estimated from the lipid-to-protein ratio used in the photo-controlled cargo release and uptake assay. For this estimate, a GUV was approximated as a sphere with a diameter ($D$) of 10 $\mu\text{m}$. After mixing the sender and receiver GUV suspensions, the final total lipid concentration ($C_{\text{lipid}}$) was 250 $\mu\text{M}$ and the concentration for each AH monomer ($C_{\text{AH}}$) was 1 $\mu\text{M}$. Thus, the nominal AH monomer-to-lipid ratio ($R$) is:

$$R = \frac{C_{\text{AH}}}{C_{\text{lipid}}} = \frac{1}{250} = 0.004$$

The membrane area of one GUV ($A_{\text{GUV}}$) was calculated as:

$$A_{\text{GUV}} = 4\pi \left(\frac{D}{2}\right)^2 = 4\pi(5)^2 = 314.16 \mu\text{m}^2$$

An average area ($a_{\text{lipid}}$) of 0.65 $\text{nm}^2$ per lipid molecule was used as an approximate value<sup>2</sup>. Because the lipid concentration represents both membrane leaflets, the total number of lipid molecules per GUV ($N_{\text{lipid,total}}$) was estimated as:

$$N_{\text{lipid,total}} = \frac{2A}{a_{\text{lipid}}} = \frac{2 \times 314.16 \times 10^6}{0.65} = 9.67 \times 10^8 (\text{copies/GUV})$$

The factor of 2 accounts for the inner and outer leaflets. Assuming that all input AH monomers insert into the outer leaflet, distribute uniformly among GUVs, and assemble completely into heptamers, the number of AH monomers ($N_{\text{AH,monomer}}$) and heptamers ($N_{\text{AH,heptamer}}$) per GUV were calculated as:

$$N_{\text{AH,monomer}} = R \times N_{\text{lipid,total}} = 0.004 \times 9.67 \times 10^8 = 3.87 \times 10^6 (\text{copies/GUV})$$

$$N_{\text{AH,heptamer}} = \frac{N_{\text{AH,monomer}}}{7} = \frac{3.87 \times 10^6}{7} = 5.52 \times 10^5 (\text{copies/GUV})$$

Under these assumptions, the nominal upper limit is therefore approximately $5.52 \times 10^5$ AH heptamers per GUV. However, the actual number of membrane-inserted AH heptamers may be lower, depending on the affinity between AH and lipids, as not all AH monomers necessarily bind to GUV membranes or assemble into heptamers. Since the actual affinity is unclear, we assume that $x\%$ ($x$, ranging from 0 to 100) of AH heptamers are successfully bound at the GUV membranes. This parameter is used in the subsequent geometric estimation (Note S2).

### Note S2. Geometric Estimation of the Capacity for AH Heptamer Accumulation at the GUV-GUV Contact Interface

We then estimated if AH heptamers could accumulate at the GUV-GUV contact area more than or similar to those at the exposed area. The GUV-GUV contact area was approximated as a flat circular disk with a contact diameter, $d$, of around 2 $\mu\text{m}$, as illustrated by the white dashed lines in Figure S14. Each AH heptamer was assumed to have a circular cross-section with a diameter of 8.5 nm (Reference 3).

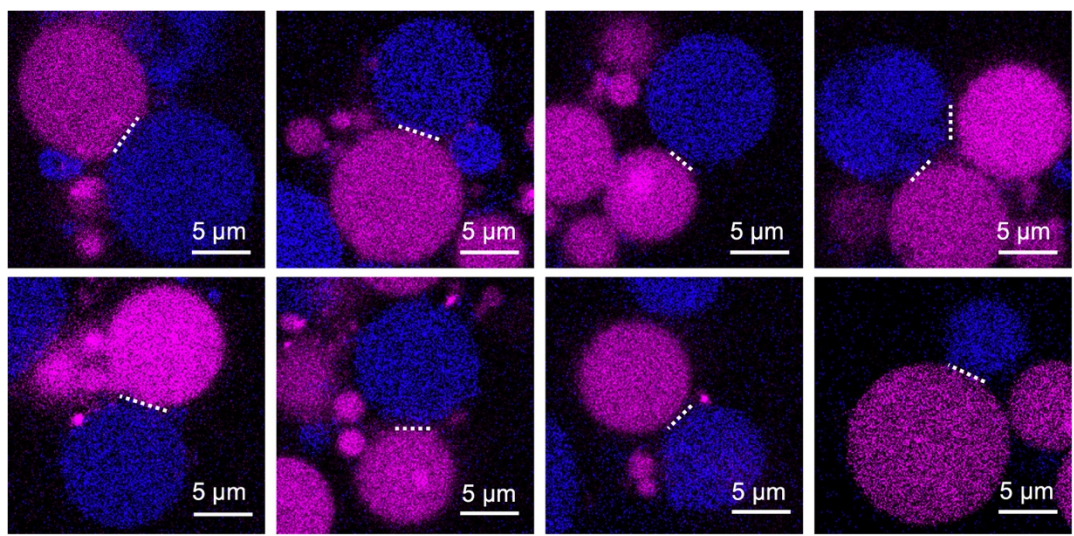

**Figure S14. Representative confocal images of linked GUV pairs selected from the photo-controlled cargo release and uptake assay.** Magenta, OA647; blue, Cascade Blue–dextran. White dashed lines indicate the apparent membrane contact regions used to define the representative contact diameter, $d$, in the geometric estimate. Scale bars: 5 $\mu\text{m}$.

The contact area ($A_{\text{contact}}$) in a flat circular dish was calculated as:

$$A_{\text{contact}} = \pi \left( \frac{d}{2} \right)^2 = 3.14 \mu\text{m}^2$$

where $d=2 \mu\text{m}$. Thus, the fraction ($f_{\text{contact}}$) of the GUV surface area (calculated in Note S1) occupied by the contact interface is:

$$f_{\text{contact}} = \frac{A_{\text{contact}}}{A_{\text{GUV}}} \times 100\% = \frac{3.14}{314} = 1\%$$

The area ($A_{\text{AH}}$) that one AH heptamer with a diameter of 8.5 nm can occupy was calculated as:

$$A_{\text{AH}} = \pi \left( \frac{8.5}{2} \right)^2 = 56.75 \text{ nm}^2$$

The maximum number of AH heptamer ($N_{\text{contact, capacity}}$) that can occupy the contact area can be estimated by assuming hexagonal close packing:

$$N_{\text{contact, capacity}} = \frac{A_{\text{contact}}}{A_{\text{AH}}} \times 0.9 = \frac{3.14 \times 10^6}{56.75} \times 0.9 \approx 4.9 \times 10^4 \text{ (copies}/A_{\text{contact}})$$

where 0.9 represents the approximated maximum two-dimensional packing fraction for identical circles arranged in a hexagonal lattice ($\pi/\sqrt{3} \approx 0.907$).

As proposed in Figure S15, ST-SC-mediated covalent linkage may promote the retention and accumulation of AH 14-mers at the contact interface. Here we assume that AH in the contact area reaches its maximum capacity, *i.e.*, $4.9 \times 10^4$ (heptamer). Under this situation and when assuming that $x\%$ of AH heptamers is bound to GUVs, the fraction of heptamers at the contact ($f_{\text{contact}}$) and exposed ($f_{\text{exposed}}$) areas are:

$$f_{\text{contact}} = 4.9 \times 10^4 / 5.52 \left( \frac{x}{100} \right) \times 10^5$$

$$f_{\text{exposed}} = 1 - 4.9 \times 10^4 / 5.52 \left( \frac{x}{100} \right) \times 10^5$$

As shown in Figure S16, the maximum fraction of membrane-inserted AH heptamers that could be accommodated at the contact area decreases as the effective membrane-incorporation fraction increases. When 10% AH heptamers are bound to GUV membranes, up to approximately 90% of heptamers will theoretically accumulate to the contact area, leaving approximately 10% on the solvent-exposed area. These results suggest that, if the actual membrane-inserted AH population is substantially lower than the nominal estimate, ST-SC-mediated recruitment and retention could potentially concentrate a large proportion of the available AH heptamers at the GUV–GUV contact interface.

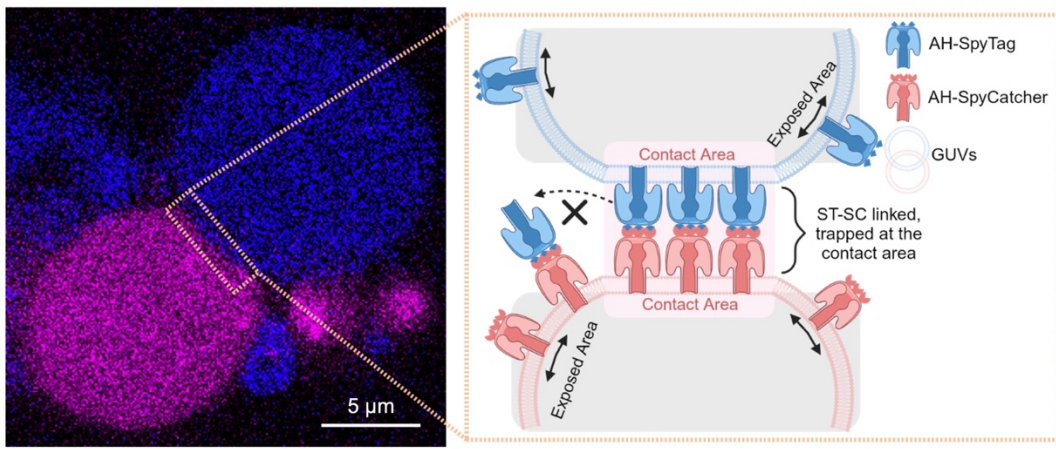

**Figure S15. Proposed mechanism of ST-SC-mediated retention of AH 14-mers at the GUV-GUV contact interface.** Left: Representative confocal fluorescence image of linked GUVs. Scale bar: 5 $\mu\text{m}$. Right: schematic illustration of the proposed

distribution and lateral mobility of AH at the GUV-GUV contact interface. The contact regions (pink shading) contain covalently linked AH 14mers, whereas the solvent-exposed membrane regions (gray shading) contain unpaired AH heptamers. Solid double-headed arrows indicate the lateral diffusion of individual AH heptamers on the exposed area on GUV membranes, whereas the dashed arrow indicates the restricted movement of covalently linked AH 14-mers at the contact interface.

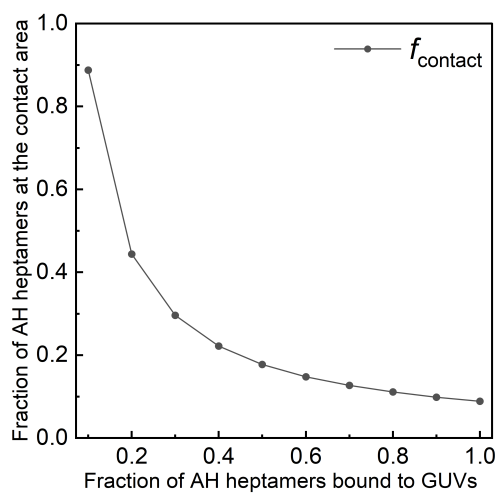

**Figure S16. Theoretical maximum fraction of membrane-inserted AH heptamers inserted at the GUV-GUV contact area.** $f_{\text{contact}}$ was calculated based on the formula shown above, which decreased with the increase of fraction that AH heptamers bound to GUV membranes. When 10% of AH heptamers are bound, around 90% of heptamers will accumulate to the contact area. These values represent theoretical upper limits rather than experimentally measured AH distributions.
